# The slowly evolving genome of the xenacoelomorph worm *Xenoturbella bocki*

**DOI:** 10.1101/2022.06.24.497508

**Authors:** Philipp H. Schiffer, Paschalis Natsidis, Daniel J. Leite, Helen E. Robertson, François Lapraz, Ferdinand Marlétaz, Bastian Fromm, Liam Baudry, Fraser Simpson, Eirik Høye, Anne-C. Zakrzewski, Paschalia Kapli, Katharina J. Hoff, Steven Mueller, Martial Marbouty, Heather Marlow, Richard R. Copley, Romain Koszul, Peter Sarkies, Maximilian J. Telford

## Abstract

The evolutionary origins of Bilateria remain enigmatic. One of the more enduring proposals highlights similarities between a cnidarian-like planula larva and simple acoel-like flatworms. This idea is based in part on the view of the Xenacoelomorpha as an outgroup to all other bilaterians which are themselves designated the Nephrozoa (protostomes and deuterostomes). Genome data can help to elucidate phylogenetic relationships and provide important comparative data. Here we assemble and analyse the genome of the simple, marine xenacoelomorph *Xenoturbella bocki*, a key species for our understanding of early bilaterian and deuterostome evolution. Our highly contiguous genome assembly of *X. bocki* has a size of ∼111 Mbp in 18 chromosome like scaffolds, with repeat content and intron, exon and intergenic space comparable to other bilaterian invertebrates. We find *X. bocki* to have a similar number of genes to other bilaterians and to have retained ancestral metazoan synteny. Key bilaterian signalling pathways are also largely complete and most bilaterian miRNAs are present. We conclude that *X. bocki* has a complex genome typical of bilaterians, in contrast to the apparent simplicity of its body plan. Overall, our data do not provide evidence supporting the idea that Xenacoelomorpha are a primitively simple outgroup to other bilaterians and gene presence/absence data support a relationship with Ambulacraria.

## Introduction

*Xenoturbella bocki* (Fig 1) is a morphologically simple marine worm first described from specimens collected from muddy sediments in the Gullmarsfjord on the West coast of Sweden. There are now 6 described species of *Xenoturbella* - the only genus in the higher-level taxon of Xenoturbellida^1^. *X. bocki* was initially included as a species within the Platyhelminthes^2^, but molecular phylogenetic studies have shown that Xenoturbellida is the sister group of the Acoelomorpha, a second clade of morphologically simple worms also originally considered Platyhelminthes: Xenoturbellida and Acoelomorpha constitute their own phylum, the Xenacoelomorpha^3,4^. The monophyly of Xenacoelomorpha is convincingly supported by their sharing unique amino acid signatures in their Caudal genes^3^ and Hox4/5/6 gene^5^. In the present work we analyse our data in this phylogenetic framework of a monophyletic taxon Xenacoelomorpha.

**Figure 1:**
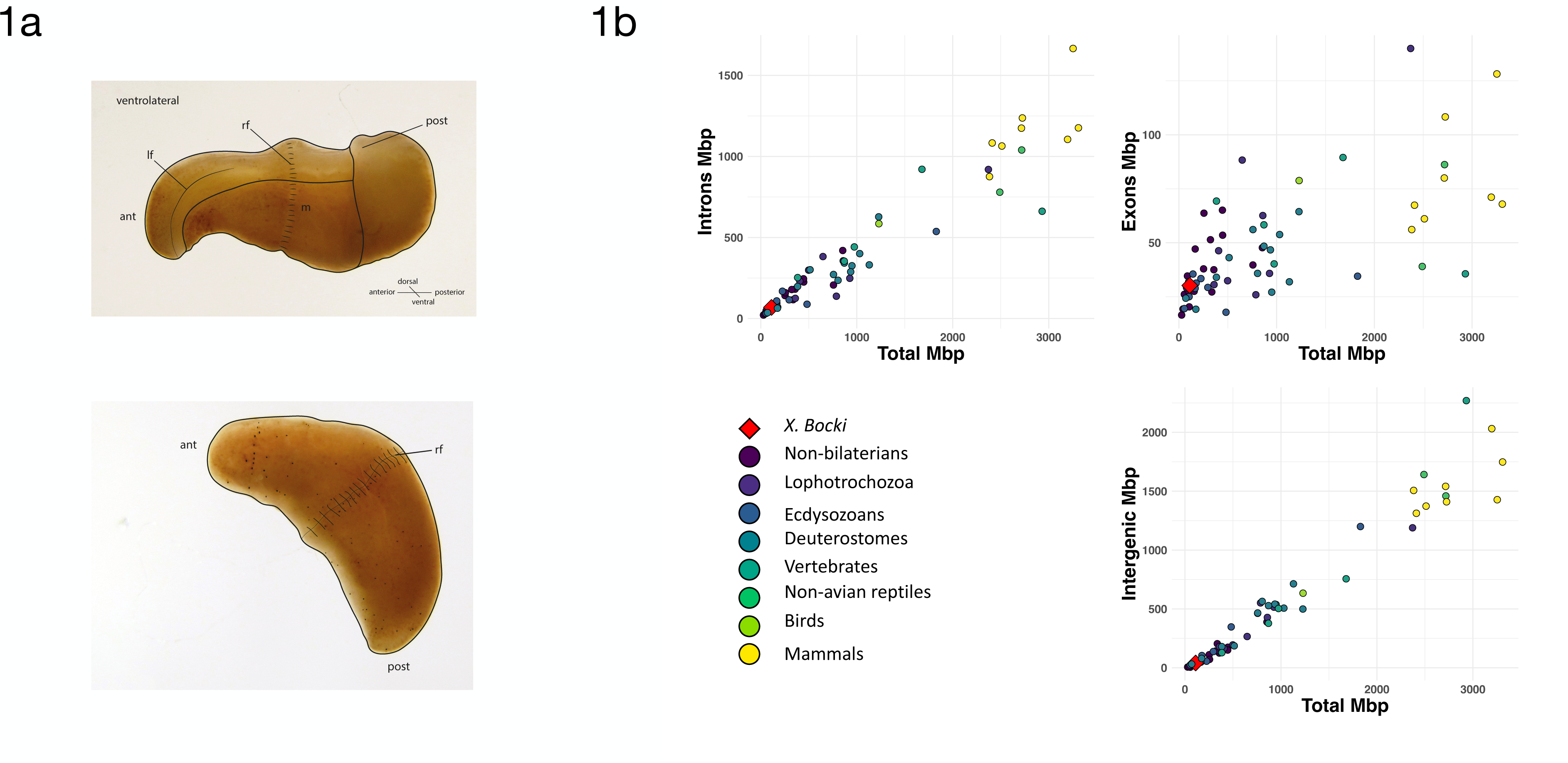
(a) Schematic drawings of *Xenoturbella bocki* showing the simple body organisation of the marine vermiform animal. Abbreviations: ant - anterior, post - posterior, lf - lateral furrow, rf - ring furrow, m - mouth opening. (b) A comparison of total length of exons, introns, and intergeneic space in the *X. bocki* genome with other metazoans (data from ref 20). *X. bocki* does not appear to be an outlier in any of theses comparisons.

The simplicity of xenacoelomorph species compared to other bilaterians is a central feature of discussions over their evolution. While Xenacoelomorpha are clearly monophyletic, their phylogenetic position within the Metazoa has been controversial for a quarter of a century. There are two broadly discussed scenarios: a majority of studies have supported a position for Xenacoelomorpha as the sister group of all other Bilateria (the Protostomia and Deuterostomia, collectively named Nephrozoa)^4,6–8;^ work we have contributed to^1,3,9,10^, has instead placed Xenacoelomorpha within the Bilateria as the sister group of the Ambulacraria (Hemichordata and Echinodermata) to form a clade called the Xenambulacraria^9^.

*Xenoturbella bocki* has neither organized gonads nor a centralized nervous system. It has a blind gut, no body cavities and lacks nephrocytes^11^. If Xenacoelomorpha is the sister group to Nephrozoa these character absences can be parsimoniously interpreted as representing the primitive state of the Bilateria. According to advocates of the Nephrozoa hypothesis, these and other characters absent in Xenacoelomorpha must have evolved in the lineage leading to Nephrozoa after the divergence of Xenacoelomorpha. More generally there has been a tendency to interpret Xenacoelomorpha (especially Acoelomorpha) as living approximations of Urbilateria^12^.

An alternative explanation for the simple body plan of xenaceolomorphs is that it is derived from that of more complex urbilaterian ancestors through loss of morphological characters. The loss or remodelling of morphological complexity is a common feature of evolution in many animal groups and is typically associated with unusual modes of living^13,14^ – in particular the adoption of a sessile (sea squirts, barnacles) or parasitic (neodermatan flatworms, orthonectids) lifestyle, extreme miniaturization (e.g. tardigrades, orthonectids), or even neoteny (e.g. flightless hexapods).

In the past some genomic features gleaned from analysis of various Xenacoelomorpha have been used to test these evolutionary hypotheses. For example, the common ancestor of the protostomes and deuterostomes has been reconstructed with approximately 8 Hox genes but only 4 have been found in the Acoelomorpha (Nemertoderma) and 5 In *Xenoturbella*. This has been interpreted as a primary absence with the full complement of 8 appearing subsequent to the divergence of Xenacoelomorpha and Nephrozoa. Similarly, analysis of the microRNAs (miRNAs) of an acoelomorph, *Symsagittifera roscoffensis*, found that many bilaterian miRNAs were absent from its genome^15^. Some of the missing bilaterian miRNAs, however, were subsequently observed in *Xenoturbella*^9^.

The few xenacoelomorph genomes available to date are from the acoel *Hofstenia miamia*^16^ – like other Acoelomorpha it shows accelerated sequence evolution relative to *Xenoturbella*^3^ – and from two closely related species *Praesagittifera naikaiensis*^17^ and *Symsagittifera roscoffensis*^18^. The analyses of gene content of *Hofstenia* showed similar numbers of genes and gene families to other bilaterians^16^, while an analysis of the neuropeptide content concluded that most bilaterian neuropeptides were present in Xenacoelomorpha^19^.

In order to infer the characteristics of the ancestral xenacoelomorph genome, and to complement the data from the Acoelomorpha, we describe a highly-scaffolded genome of the slowly evolving xenacoelomorph *Xenoturbella bocki*. This allows us to contribute knowledge of Xenacoelomorpha and *Xenoturbella* in particular of genomic traits, such as gene content and genome-structure and to help reconstruct the genome structure and composition of the ancestral xenacoelomorph.

## Results

### Assembly of a draft genome of *Xenoturbella bocki*

We collected *Xenoturbella bocki* specimens (Fig. 1) from the bottom of the Gullmarsfjord close to the biological field station in Kristineberg (Sweden). These adult specimens were starved for several days in tubes with artificial sea water, and then sacrificed in lysis buffer. We extracted high molecular weight (HMW) DNA from single individuals for each of the different sequencing steps below.

We assembled a high-quality draft genome of *Xenoturbella bocki* using one short read Illumina library and one TruSeq Synthetic Long Reads (TSLR) Illumina library. We used a workflow based on a primary assembly with SPAdes (Methods; ^20^). The primary assembly had an N50 of 8.5kb over 37,880 contigs with a maximum length of 206,709bp. After using the redundans pipeline^21^ this increased to an N50 of ∼62kb over 23,094 contigs and scaffolds spanning ∼121Mb, and a longest scaffold of 960,978kb (supplementary Table 1).

The final genome was obtained with Hi-C scaffolding using the program instaGRAAL (Methods, see supplementary for contact map;^22^). The scaffolded genome has a span of 111 Mbp (117 Mbp including small fragments unincorporated into the HiC assembly) and an N50 of 2.7 Mbp (for contigs >500bp). The assembly contains 18 megabase-scale scaffolds encompassing 72 Mbp (62%) of the genomic sequence, with 43% GC content. The original assembly indicated a repeat content of about 25% after a RepeatModeller based RepeatMasker annotation (Methods). As often seen in non-model organisms, about 2/3 of the repeats are not classified.

We used BRAKER1^23,24^ with extensive RNA-Seq data, and additional single-cell UTR enriched transcriptome sequencing data to predict 15,154 gene models. 9,575 gene models (63%) are found on the 18 large scaffolds (which represent 62% of the total sequence). 13,298 of our predicted genes (88%) have RNA-Seq support. Although this proportion is at the low end of bilaterian gene counts, we note that our RNA-seq libraries were all taken from adult animals and thus may not represent the true complexity of the gene complement. We consider our predicted gene number to be a lower bound estimate for the true gene content.

The predicted *X. bocki* genes have a median coding length of 873 nt and a mean length of 1330 nt. Median exon length is 132 nt (mean 212 nt) and median intron length is 131 nt (mean 394 nt). Genes have a median of 4 exons and a mean of 8.5 exons. 2,532 genes have a single exon and, of these, 1,381 are supported as having a single exon by RNA-Seq (TPM>1). A comparison of the exon, intron, and intergenic sequence content in *Xenoturbella* with those described in other animal genomes^25^ show that *X. bocki* falls within the range of other similarly sized metazoan genomes (Fig. 1b) for all these measures.

### The genome of a co-sequenced *Chlamydia* species

We recovered the genome of a marine *Chlamydia* species from Illumina data obtained from one *X. bocki* specimen and from Oxford Nanopore data from a second specimen supporting previous microscopic analyses and single gene PCRs suggesting that *X. bocki* is host to a species in the bacterial genus *Chlamydia*. The bacterial genome was found as 5 contigs spanning 1,906,303 bp (N50 of 1,237,287 bp) which were assembled into 2 large scaffolds. Using PROKKA^26^, we predicted 1,738 genes in this bacterial genome, with 3 ribosomal RNAs, 35 transfer RNAs, and 1 transfer-messenger RNA. The genome is 97.5% complete for bacterial BUSCO^27^ genes, missing only one of the 40 core genes.

Marine chlamydiae are not closely related to the group of human pathogens^28^ and we were not able to align the genome of the *Chlamydia*-related symbioint from *X. bocki* to the reference strain *Chlamydia trachomatis* F/SW4, nor to *Chlamydophila pneumoniae* TW-183. To investigate the phylogenetic position of the species co-occurring with *Xenoturbella*, we aligned the 16S rRNA gene from the *X. bocki*-hosted *Chlamydia* with orthologs from related species including sequences of genes amplified from DNA/RNA extracted from deep sea sediments. The *X. bocki*-hosted *Chlamydia* belong to a group designated as Simkaniaceae in^28^, with the sister taxon in our phylogenetic tree being the *Chlamydia* species previously found in *X. westbladi* (*X. westbladi* is almost certainly a synonym of *X. bocki*)^7^ (Fig. 2a).

**Figure 2a:**
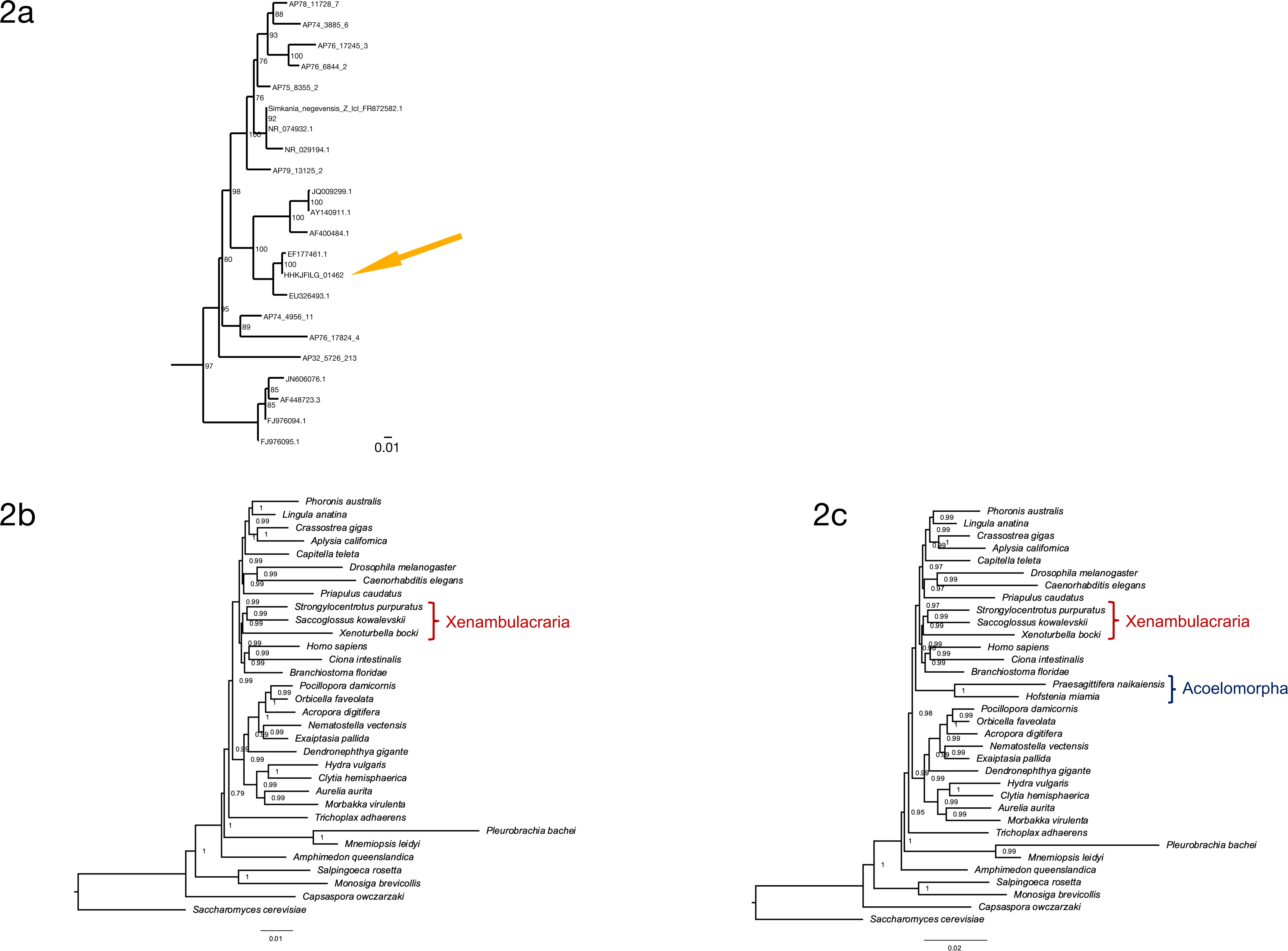
*Xenoturbella bocki* harbours a marine Chlamydiae species as potential symbiont. In the phylogenetic analysis of 16 S r DNA (ML: GTR+F+R7; bootstrap values included) the bacteria in our *X. bocki* isolate (arrow) are sister to a previous isolate from *X. westbaldi*. *X. westbaldi* is most likely a mis-identification of *X. bocki.* (b/c) A phylogeny based on presence and absence of genes calculated with OMA. Both analysis (b) and (c) confirm Xenambularcraria, i.e. Xenoturbellida in a group with Echinoderms and Hemichordates. Inclusion of the acoel flat worms places these as sister to all other Bilateria (b). This placement appears an artefact due to the very fast evolution in this taxon, in particular as good evidence exists for uniting Xenoturbellida and Acoela refs 5 and 6.

To investigate whether the *X. bocki*-hosted *Chlamydia* might contribute to the metabolic pathways of its host, we compared the completeness of metabolic pathways in KEGG for the *X. bocki* genome alone and for the *X. bocki* genome in combination with the bacteria. We found only slightly higher completeness in a small number of pathways involved in carbohydrate metabolism, carbon fixation, and amino acid metabolism (see supplementary material) suggesting that the relationship is likely to be commensal or parasitic rather than a true symbiosis.

A second large fraction of bacterial reads, annotated as Gammaproteobacteria, were identified and filtered out during the data processing steps. These bacteria were also previously reported as potential symbionts of *X. bocki*^29^. However, these sequences were not sufficiently well covered to reconstruct a genome and we did not investigate them further.

### A phylogenetic gene presence/absence matrix supports Xenambulacraria

The general completeness of the *X. bocki* gene set allowed us to use the presence and absence of genes identified in our genomes as a source of information to find the best supported phylogenetic position of the Xenacoelomorpha. We conducted two separate phylogenetic analyses of gene presence/absence data: one including the fast-evolving Acoelomorpha and one without. In both analyses the best tree grouped *Xenoturbella* with the Ambulacraria (Fig. 2b). The analysis including acoels, however, placed the acoels as the sister-group to Nephrozoa separate from *Xenoturbella* (Fig. 2c). Because other data have shown the monophyly of Xenacoelomorpha to be robust, we interpret this result as being the result of systematic error caused by a high rate of gene loss or by orthologs being incorrectly scored as missing due to higher rates of sequence evolution in acoelomorphs^30^.

### The *X. bocki* molecular toolkit is typical of bilaterians

One of our principal aims was to ask whether the *Xenoturbella* genome lacks characteristics otherwise present in the Bilateria. We found that for the Metazoa gene set in BUSCO (v5) the *X. bocki* proteome translated from our gene predictions is 82.5% complete and ∼90% complete when partial hits are included (82% and 93% respectively for the Eukaryote gene set). This estimate is even higher in the acoel *Hofstenia miamia*, which was originally reported to be 90%^16^, but in our re-analysis was 95.71%. In comparison, the morphologically highly simplified and fast evolving annelid *Intoshia linei*^31^ has a genome of fewer than 10,000 genes^32^ and in our analysis is only ∼64% complete for the BUSCO (v5) Metazoa set. The model nematode *Caenorhabditis elegans* is ∼79% complete for the same set. Despite the morphological simplicity of both *Xenoturbella*, and *Hofstenia,* these Xenacoelomorpha are missing few core genes compared to other bilaterian lineages that we perceive to have undergone a high degree of morphological evolutionary change (such as the evolution of miniaturisation, parasitism, sessility etc).

Using our phylogenomic matrix of gene presence/absence (see above) we identified all orthologs that could be detected both in Bilateria (in any bilaterian lineage) and in any non-bilaterian; ignoring horizontal gene transfer and other rare events, these genes must have existed in Urbilateria (and, of less interest to us, in Urmetazoa). The absence of any of these bilaterian genes in any lineage of Bilateria must therefore be explained by loss of the gene. All individual bilaterian genomes were missing many of these orthologs but Xenacoelomorphs and some other bilaterians lacked more of these than did other taxa. The average numbers of these genes present in bilaterians = 7577; *Xenoturbella* = 5459; *Hofstenia* = 5438; *Praesagittifera* = 4280; *Drosophila* = 4844; *Caenorhabditis* = 4323.

To better profile the *Xenoturbella* and xenacoelomorph molecular toolkit, we used OrthoFinder to conduct orthology searches in a comparison of 155 metazoan and outgroup species, including the transcriptomes of the sister species *X. profunda* and an early draft genome of the acoel *Paratomella rubra* we had available, as well as the *Hofstenia* and *Praesagittifera* proteomes (Supplementary online material). For each species we counted, in each of the three Xenacoelomorphs, the number of orthogroups for which a gene was present. The proportion of orthogroups containing an *X. bocki* and *X. profunda* protein (87.4% and 89.2%) are broadly similar to the proportions seen in other well characterised genomes, for example *S. purpuratus* proteins (93.8%) or *N. vectensis* proteins (84.3%) (Fig 3a). In this analysis, the fast-evolving nematode *Caenorhabditis elegans* appears as an outlier, with only ∼64% of its proteins in orthogroups and ∼35% unassigned. Both *Xenoturbella* species have an intermediate number of unassigned genes of ∼11-12%. Similarly, the proportion of species-specific genes (∼14% of all genes) corresponds closely to what is seen in most other species (with the exception of the parasitic annelid *I. linei*, Fig. 3a).

**Figure 3:**
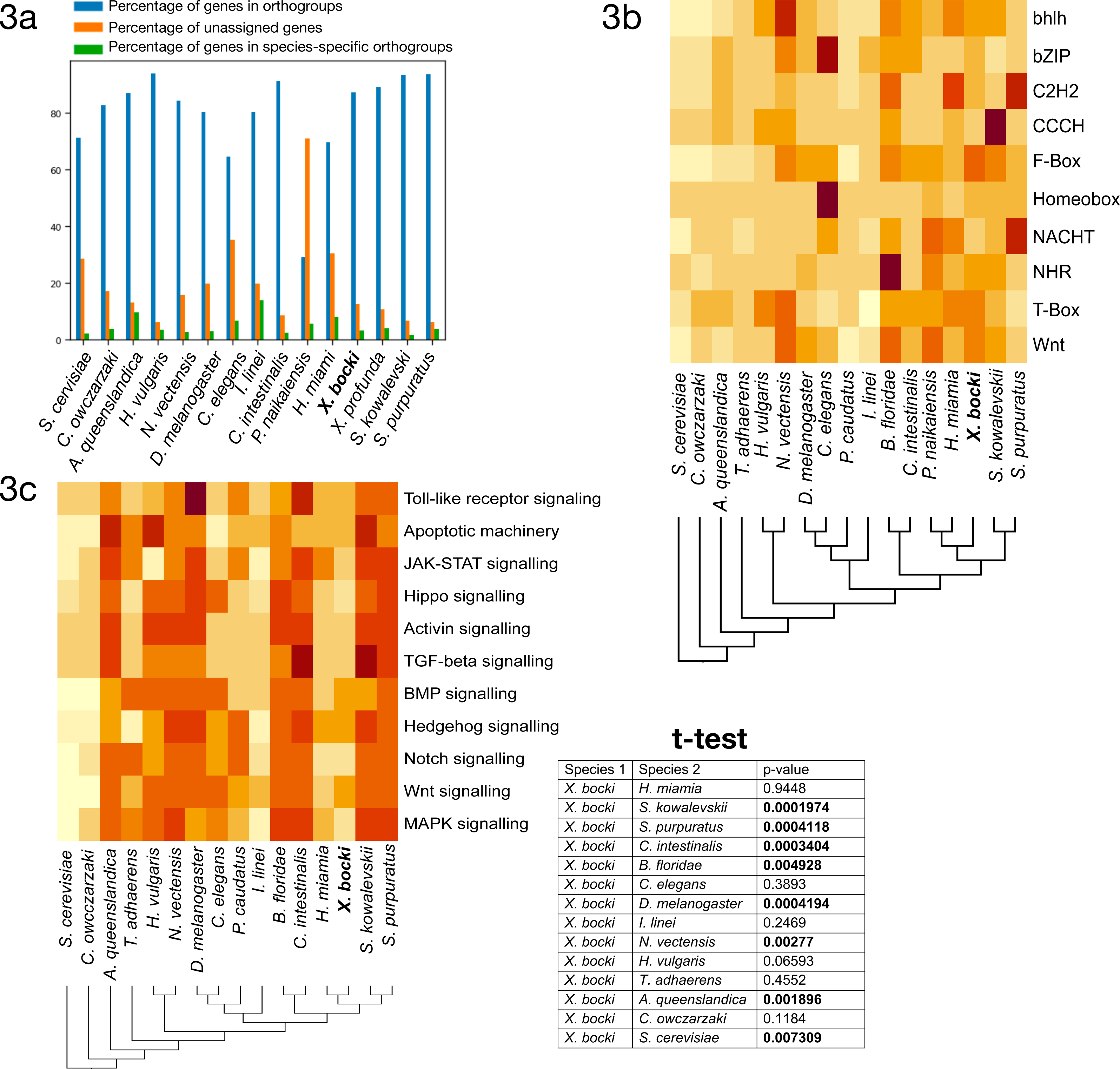
(a) In our orthology screen *X. bocki* shows similar percentages of genes in orthogroups, unassigned genes, and species-specific orthogroups as other well-annotated genomes. (b) The number of family members per species in major gene families (based on Pfam domains), like transcription factors, fluctuates in evolution. The *X. bocki* genome does not appear to contain particularly less or more genes in any of the analysed families. (c) Cell signalling pathways in *X. bocki* are functionally complete, but in comparison to other species contain less genes. The overall completeness is not significantly different to, for example, the nematode *C. elegans* (inset, t-test). Schematic cladograms in b/c drawn by the authors.

### Idiosyncrasies of *Xenoturbella*

In order to identify sets of orthologs specific to the two *Xenoturbella* species we used the kinfin software^33^ and found 867 such groups in the OrthoFinder clustering. We profiled these genes based on Pfam domains and GO terms derived from InterProScan. While these *Xenoturbella* specific proteins fall into diverse classes, we did see a considerable number of C-type lectin, Immunoglobulin-like, PAN, and Kringle domain containing Pfam annotations. Along with the Cysteine-rich secretory protein family and the G-protein coupled receptor activity GO terms, these genes and families of genes may be interesting for future studies into the biology of *Xenoturbella* in its native environment.

### Gene families and signaling pathways are retained in *X. bocki*

In our orthology clustering we did not see an inflation of *Xenoturbella*-specific groups in comparison to other taxa, but also no conspicuous absence of major gene families (Fig. 3b). Family numbers of transcription factors like Zinc-fingers or homeobox-containing genes, as well as, for example, NACHT-domain encoding genes seem to be neither drastically inflated nor contracted in comparison to other species in our InterProScan based analysis.

To catalogue the completeness of cell signalling pathways we screened the *X. bocki* proteome against KEGG pathway maps using GenomeMaple^34^. The *X. bocki* gene set is largely complete in regard to the core proteins of these pathways, while an array of effector proteins is absent (Fig. 3c). In comparison to other metazoan species, as well as a unicellular choanoflagellate and a yeast, the *X. bocki* molecular toolkit has significantly lower KEGG completeness than morphologically complex animals such as the sea urchin and amphioxus (t-test; Fig. 3c). *Xenoturbella* is, however, not significantly less complete when compared to other bilaterians considered to have low morphological complexity and which have been shown to have reduced gene content, such as *C. elegans*, the annelid parasite *Intoshia linei*, or the acoel *Hofstenia miamia* (Fig. 3c).

### Clustered homeobox genes in the *X. bocki* genome

Acoelomorph flatworms possess three unlinked HOX genes, orthologs of anterior (Hox1), central (Hox4/5 or Hox5) and posterior Hox (HoxP). In contrast, previous analysis of *X. bocki* transcriptomes identified one anterior, three central and one posterior Hox genes. We identified clear evidence of a syntenic Hox cluster with four Hox genes (centHox1, postHox, centHox3, and antHox1) in the *X. bocki* genome (Fig. 4). There was also evidence of a fragmented annotation of centHox2, split between the 4 gene Hox cluster and a separate scaffold (Fig. 4). In summary, this suggests that all five Hox genes form a Hox cluster in the *X. bocki* genome, but that there are possible unresolved assembly errors disrupting the current annotation. We also identified other homeobox genes on the Hox cluster scaffold, including Evx (Fig. 4a). Along with the Hox genes, we surveyed other homeobox genes that are typically clustered in Bilateria. The canonical bilaterian paraHox cluster contains three genes Cdx, Xlox (=Pdx) and Gsx. We identified Cdx and a new Gsx annotation on the same scaffold, as well as a previously reported Gsx paralog on a separate scaffold. This indicates partial retention of the paraHox cluster in *X. bocki* along with a duplication of Gsx. On both of these paraHox containing scaffolds we observed other homeobox genes.

**Figure 4:**
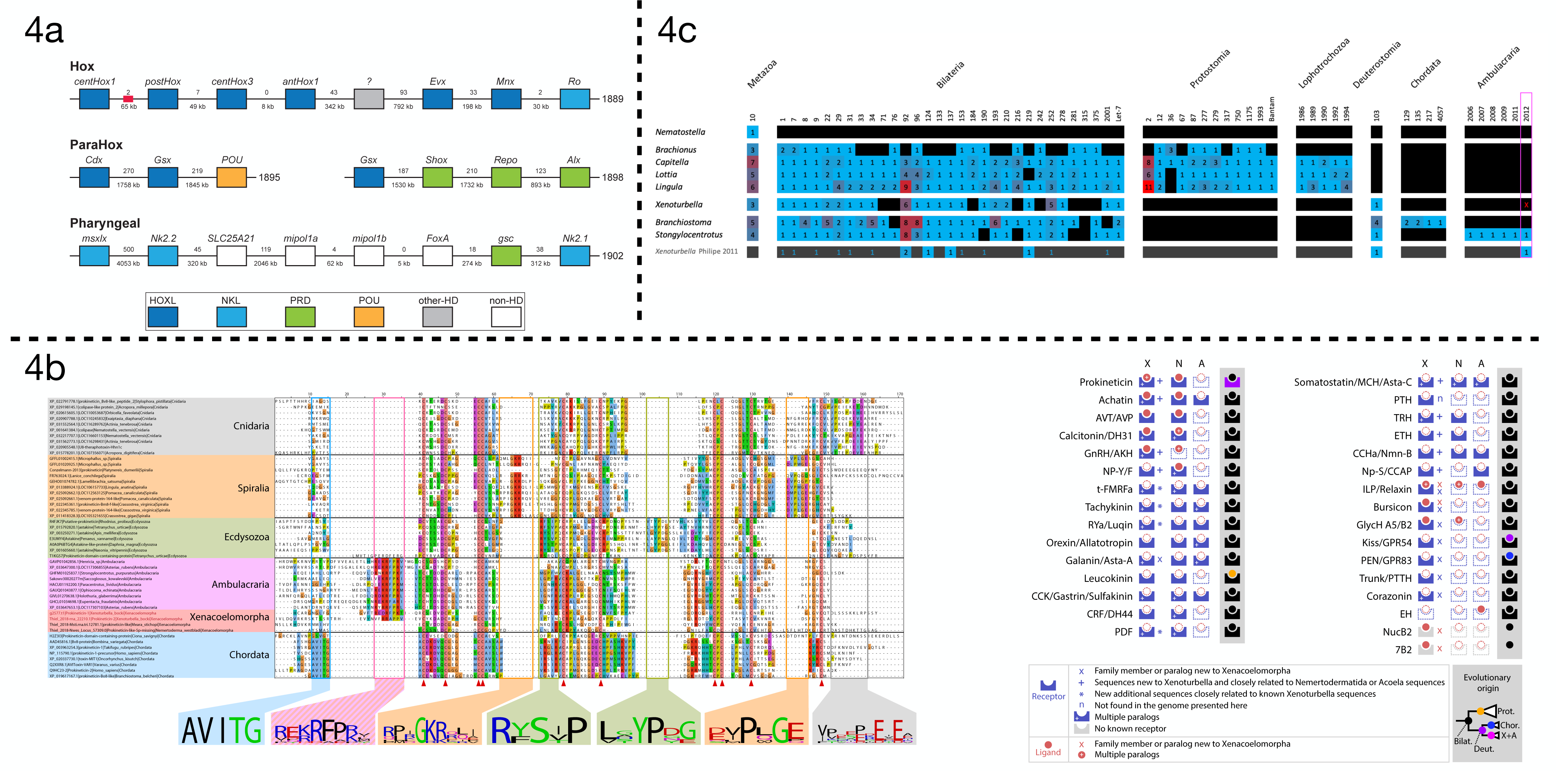
(a) *X. bocki* has 5 HOX genes, which are located in relatively close proximity on one of our chromosome size scaffolds. Similar clusters exist for the ParaHox and “pharyngeal” genes. Numbers between genes are distance (below) and number of genes between (below). Colours indicate gene families. Red box marks the position of a partial Hox gene. The “?” gene has an unresolved homeodomain identity. (b) We found a specific prokineticin ligand signature sequence in *X. bocki*, which was previously reported for Ecdysozoa and Chordata, as well as a “K/R-RFP-K/R”, sequence shared only by ambulacrarians and *X. bocki*. Signature previously reported for Ecdysozoa and Chordata, as well as new signatures we found in Spiralia and Cnidaria is absent from ambulacrarian and *X. bocki* prokineticin ligand sequences. The inset cladogram depicts the evolutionary origin of sequences in accordance with our analysis: **Bilat**erian, **Prot**ostomia, **Chor**date, **X**enacoelomorpha + **A**mbulacraria last common ancestor respectively. (c) The revised microRNA complement of *X. bocki* has a near complete set of metazoan, bilaterian and deuterostome families and genes. Presence (color) and absence (black) of microRNA families (column), paralogue numbers (values & heatmap coloring) organized in node-specific blocks in a range of representative protostome and deuterostome species compared with *Xenoturbella* (species from MirGeneDB 2.1 - Fromm et al 2021). The bottom row depicts 2011 complement by Philippe et al 2011 (blue numbers on black depict detected miRNA reads, but lack of genomic evidence). Red “x” in pink box highlights the lack of evidence for an ambulacraria-specific microRNA in *X. bocki*.

Hemichordates and chordates have a conserved cluster of genes involved in patterning their pharyngeal pores - the so-called ‘pharyngeal cluster’. The homeobox genes of this cluster (Msxlx, Nk2-1/2/4/8) were present on a single *X. bocki* scaffold. Another pharyngeal cluster transcription factor, the Forkhead containing Foxa, and ‘bystander’ genes from that cluster including Egln, Mipol1 and Slc25a21 are found in the same genomic region. Different sub-parts of the cluster are found in non-bilaterians and protostomes and the cluster may well be plesiomorphic for the Bilateria rather than a deuterostome synapomorphy^35^.

### The *X. bocki* neuropeptide complement is larger than previously thought

A catalogue of acoelomorph neuropeptides was previously described using transcriptome data^36^. We have discovered 12 additional neuropeptide genes and 39 new neuropeptide receptors in *X. bocki* adding 6 bilaterian peptidergic systems to the *Xenoturbella* catalogue (NPY-F; MCH/Asta-C; TRH; ETH; CCHa/Nmn-B; Np-S/CCAP), and 6 additional bilaterian systems to the Xenacoelomorpha catalogue (Corazonin; Kiss/GPR54; GPR83; 7B2; Trunk/PTTH; NUCB2) making a total of 31 peptidergic systems (Fig. 4, Supplementary).

Among the ligand genes, we identified 6 new repeat-containing sequences. One of these, the LRIGamide-peptide, had been identified in Nemertodermatida and Acoela and its loss in *Xenoturbella* had been proposed^36^. We also identified the first 7B2 neuropeptide and NucB2/Nesfatin genes in Xenacoelomorpha. Finally, we identified 3 new *X. bocki* insulin-like peptides, one of them sharing sequence similarity and an atypical cysteine pattern with the Ambulacrarian octinsulin, constituting a potential synapomorphy of Xenambulacraria (see Supplementary).

Our searches also revealed the presence of components of the arthropod moulting pathway components (PTTH/trunk, NP-S/CCAP and Bursicon receptors), which have recently been shown to be of ancient origin (de Oliveira et al., 2019). We further identified multiple paralogs for, e.g the Tachykinin, Rya/Luquin, tFMRFa, Corazonin, Achatin, CCK, and Prokineticin receptor families. Two complete *X. bocki* Prokineticin ligands were also found in our survey (see Supplementary).

Chordate Prokineticin ligands possess a conserved N-terminal “AVIT” sequence required for the receptor activation^37^. This sequence is absent in arthropod Astakine, which instead possess two signature sequences within their Prokineticin domain ^38^. To investigate Prokineticin ligands in Xenacoelomorpha we compared the sequences of their prokineticin ligands with those of other bilaterians (Fig. 4b, Supplementary). Our alignment reveals clade specific signatures already reported in Ecdysozoa and Chordata sequences, but also two new signatures specific to Lophotrochozoa and Cnidaria sequences, as well as a very specific “K/R-RFP-K/R” signature shared only by ambulacrarian and *Xenoturbella bocki* sequences. The shared Ambulacrarian/Xenacoelomorpha signature is found at the same position as the Chordate sequence involved in receptor activation - adjacent to the N-terminus of the Prokineticin domain (Fig. 4b).

### The *X. bocki* genome contains most bilaterian miRNAs reported missing from acoels

microRNAs have previously been used to investigate the phylogenetic position of the acoels and *Xenoturbella.* The acoel *Symsagittifera roscoffensis* lacks protostome and bilaterian miRNAs and this lack was interpreted as supporting the position of acoels as sister-group to the Nephrozoa. Based on shallow 454 microRNA sequencing (and sparse genomic traces) of *Xenoturbella*, some of the bilaterian miRNAs missing from acoels were found - 16 of the 32 expected metazoan (1 miRNA) and bilaterian (31 miRNAs) microRNA families – of which 6 could be identified in genome traces^9^.

By deep sequencing two independent small RNA samples, we have now identified the majority of the missing metazoan and bilaterian microRNAs and identified them in the genome assembly (Fig. 4c). Altogether, we found 23 out of 31 bilaterian microRNA families (35 genes including duplicates); the single known Metazoan microRNA family (MIR-10) in 2 copies; the Deuterostome-specific MIR-103; and 7 *Xenoturbella*-specific microRNAs giving a total of 46 microRNA genes. None of the protostome-specific miRNAs were found. We could not confirm in the RNA sequences or new assembly a previously identified, and supposedly xenambulacrarian-specific MIR-2012 ortholog.

### The *X. bocki* genome retains ancestral metazoan linkage groups

The availability of chromosome-scale genomes has made it possible to reconstruct 24 ancestral linkage units broadly preserved in bilaterians^39^. In fast-evolving genomes, such as those of nematodes, tunicates or platyhelminths, these ancestral linkage groups (ALGs) are often dispersed and/or extensively fused (Supplementary). We were interested to test if the general conservation of the gene content in *X. bocki* is reflected in its genome structure.

We compared the genome of *Xenoturbella* to several other metazoan genomes and found that it has retained most of these ancestral bilaterian units: 12 chromosomes in the *X. bocki* genome derive from a single ALG, five chromosomes are made of the fusion of two ALGs, and one *Xenoturbella* chromosome is a fusion of three ALGs, as highlighted with the comparison of ortholog content with amphioxus, the sea urchin and the sea scallop (Fig. 5 and Supplementary).

**Figure 5:**
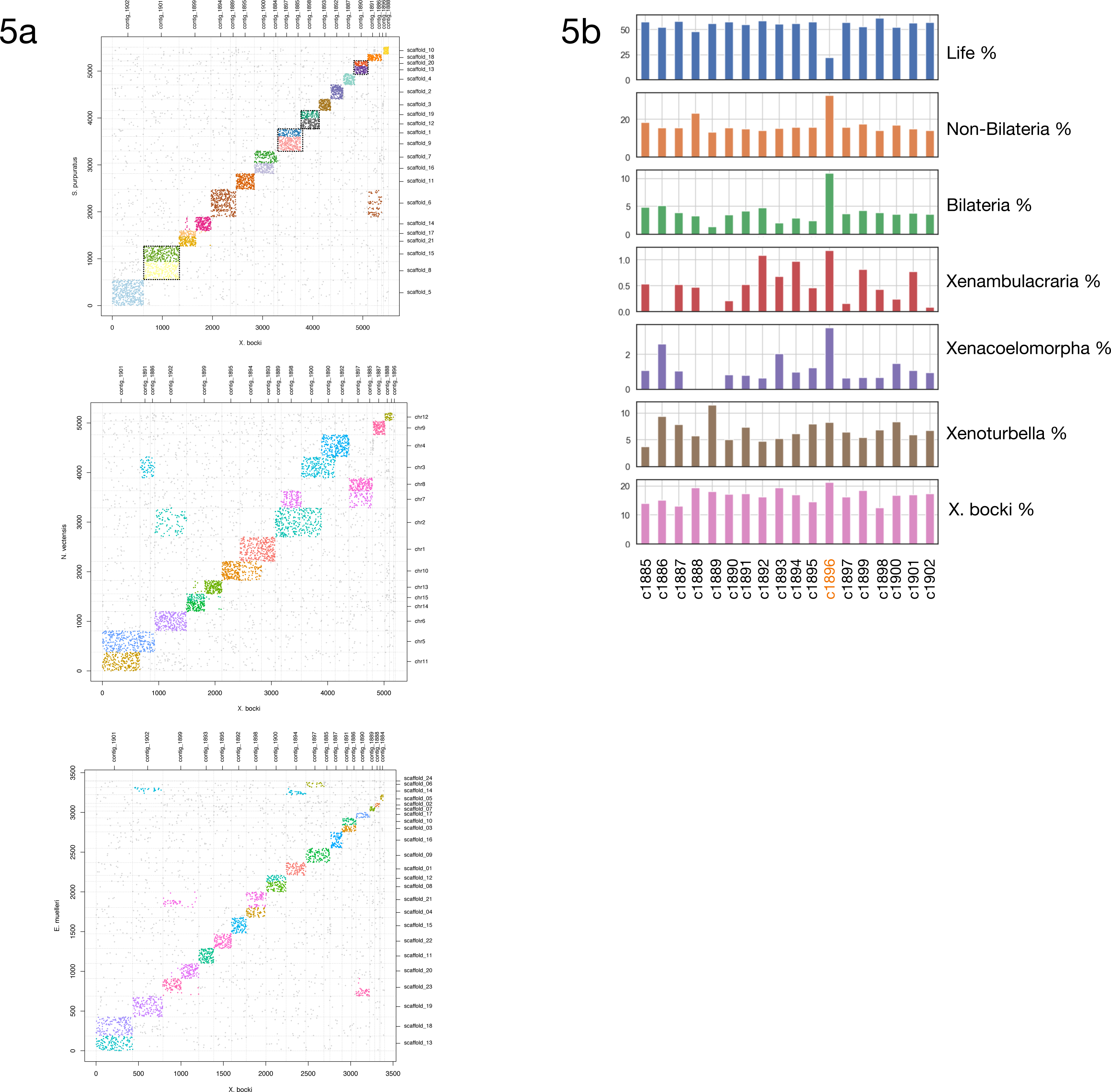
(a) A comparison of scaffolds in the *X. bocki* genome with other Metazoa. 17 of the 18 large scaffolds in the *X. bocki* genome are linked via synteny to distinct chromosomal scaffolds in these species. (b) Phylostratigraphic age distribution of genes on all major scaffolds in the *X. bocki* genome. One scaffold (c1896), which showed no synteny to a distinct chromosomal scaffold in the other metazoan species also had a divergent gene age structure in comparison to other *X. bocki* scaffolds.

One ancestral linkage group that has been lost in chordates but not in ambulacrarians nor in molluscs (ALG R in sea urchin and sea scallop) is detectable in *X. bocki* (Fig. 5), while *X. bocki* does not show the fusions that are characteristic of lophotrochozoans.

We also attempted to detect some pre-bilaterian arrangement of ancestral linkage: for instance, ref ^40^ predicted that several pre-bilaterian linkage groups successively fused in the bilaterian lineage to give ALGs A1, Q and E. These ALGs are all represented as single units in *X. bocki* in common with other Bilateria. None of the inferred pre-bilaterian chromosomal arrangements that could have provided support for the Nephrozoa hypothesis were found *in X. bocki* although of course this does not rule out Nephrozoa.

### One *X. bocki* chromosomal fragment appears aberrant

The smallest of the 18 large scaffolds in the *X. bocki* genome did not show strong 1:1 clustering with any scaffold/chromosome of the bilaterian species we compared it to. To exclude potential contamination in the assembly as a source for this contig we examined the orthogroups to which the genes from this scaffold belong. We found that *Xenoturbella profunda*^41^, for which a transcriptome is available, was the species that most often occurred in the same orthogroup with genes from this scaffold (41 shared orthogroups), suggesting the scaffold is not a contaminant.

We did observe links between the aberrant scaffold and several scaffolds from the genome of the sponge *E. muelleri* in regard to synteny, but could not detect distinct synteny relationships to a single scaffold in another species. In line with this, genes on the scaffold show a different age structure compared to other scaffolds, with both more older genes (pre bilaterian) and more *Xenoturbella* specific genes (Fig. 5b; supported by Ks statistics, Supplementary). This aberrant scaffold also had significantly lower levels of methylation than the rest of the genome (Supplementary).

## DISCUSSION

The phylogenetic positions of *Xenoturbella* and the Acoelomorpha have been controversial since the first molecular data from these species appeared over twenty five years ago. Today we understand that they constitute a monophyletic group of morphologically simple worms^1,9,42^, but there remains a disagreement over whether they represent a secondarily simplified sister group of the Ambulacraria or a primitively simple sister group to all other Bilateria.

Previous analyses of the content of genomes, especially of Acoela, have been used to bolster the latter view, with the small number of Hox genes and of microRNAs of acoels being interpreted as representing an intermediate stage on the path to the ∼8 Hox genes and 30 odd microRNAs of the Nephrozoa. A strong version of the Nephrozoa idea would go further than these examples and anticipate, for example, a genome-wide paucity of bilaterian genes, GRNs and biochemical pathways and/or an arrangement of chromosomal segments intermediate between those of the Eumetazoa and the Nephrozoa.

One criticism of the results from analyses of acoel genomes is that the Acoelomorpha have evolved rapidly (their long branches in phylogenetic trees showing high rates of sequence change). This rapid evolution might plausibly be expected to correlate with other aspects of rapid genome evolution such as higher rates of gene loss and chromosomal rearrangements leading to significant differences from other Bilateria. The more normal rates of sequence evolution observed in *Xenoturbella* therefore recommend it as a more appropriate xenacoelomorph to study with fewer apomorphic characters expected.

We have sequenced, assembled, and analysed a draft genome of *Xenoturbella bocki*. To help with annotation of the genome we have also sequenced miRNAs and small RNAs as well as using bisulphite sequencing, Hi-C and Oxford nanopore. We compared the gene content of the *Xenoturbella* genome to species across the Metazoa and its genome structure to several other high-quality draft animal genomes.

We found the *X. bocki* genome to be fairly compact, but not unusually reduced in size compared to many other bilaterians. It appears to contain a similar number of genes (∼15,000) as other animals, for example from the model organisms *D. melanogaster* (>14,000) and *C. elegans (*∼20,000). The BUSCO completeness, as well as a high level of representation of *X. bocki* proteins in the orthogroups of our 155 species orthology screen indicates that we have annotated a near complete gene set. Surprisingly, there are fewer genes than in the acoel *Hofstenia* (>22,000; BUSCO_v5 score ∼95%). This said, of the genes found in Urbilateria (orthogroups in our presence/absence analysis containing a member from both a bilaterian and an outgroup) *Xenoturbella* and *Hofstenia* have very similar numbers (5459 and 5438 respectively). Gene, intron and exon lengths all also fall within the range seen in many other invertebrate species^25^. It thus seems that basic genomic features in *Xenoturbella* are not anomalous among Bilateria. Unlike some extremely simplified animals, such as orthonectids, we observe no extreme reduction in gene content.

All classes of homeodomain transcription factors have previously been reported to exist in Xenacoelomorpha^43^. We have identified 5 HOX-genes in *X. bocki* and at least four, and probably all five of these are on one chromosomal scaffold within 187 Kbp. *X. bocki* also has the parahox genes Gsx and Cdx; while Xlox/pdx is not found, it is present in Cnidarians and must therefore have been lost^44^. If block duplication models of Hox and Parahox evolutionary relationships are correct, the presence of a complete set of parahox genes implies the existence of their Hox paralogs in the ancestor of Xenacoelomorphs suggesting the xenacoelomorph ancestor also possessed a Hox 3 ortholog. If anthozoans also have an ortholog of bilaterian Hox 2^45^, this must also have been lost from Xenacoelomorphs. The minimal number of Hox genes in the xenacoelomorph stem lineage was therefore probably 7 (AntHox1, lost Hox2, lost Hox 3, CentHox 1, CentHox 2, CentHox 3 and postHoxP).

Based on early sequencing technology and without a reference genome available, it was thought that Acoelomorpha lack many bilaterian microRNAs. Using deep sequencing of small RNAs and our high-quality genome, we have shown that *Xenoturbella* shows a near-complete bilaterian set of miRNAs including the single deuterostome-specific miRNA family (MIR-103) (Figure X). The low number of differential family losses of *Xenoturbella* (8 of 31 bilaterian miRNA families) inferred is the same as the number lost in the flatworm *Schmidtea,* and substantially lower than the number lost in the rotifer *Brachionus* (which has lost 14 bilaterian families). It is worth mentioning that *X. bocki* shares the absence of one miRNA family (MIR-216) with all Ambulacrarians although if Deuterostomia are paraphyletic this could be interpretable as a primitive state^35^.

The last decade has seen a re-evaluation of our understanding of the evolution of the neuropeptide signaling genes^46,47^. The peptidergic systems are thought to have undergone a diversification that produced approximately 30 systems in the bilaterian common ancestor^46,47^. Our study identified 31 neuropeptide systems in *X.bocki* and for all of these either the ligand, receptor, or both are also present in both protostomes and deuterostomes indicating conservation across Bilateria. It is likely that more ligands (which are short and variable) remain to be found with better detection methods. It appears that the *Xenoturbella* genome contains a nearly complete bilaterian neuropeptide complement with no signs of simplification but rather signs of expansions of certain gene families. Our analyses also reveal a potential synapomorphy linking Xenacoelomorpha with Ambulacraria (Fig 4 and Supplementary).

We have used the predicted presence and absence of genes across a selection of metazoan genomes as characters for phylogenetic analyses. Our trees re-confirm the findings of recent phylogenomic gene alignment studies in linking *Xenoturbella* to the Ambulacraria. We also used these data to test different bilaterians for their propensity to lose otherwise conserved genes (or for our inability to identify orthologs^30^). While the degree of gene loss appears similar between *Xenoturbella* and acoels, the phylogenetic analysis shows longer branches leading to the acoels, most likely due to faster evolution, gain of lineages specific genes, and some degree of gene loss in the branch leading to the Acoelomorpha. Recent work has shown the tendency of rapidly evolving genes (such as those belonging to rapidly evolving species) to be missed by orthology detection software^48,49^.

This pattern of conservation of evolutionarily old parts of the Metazoan genome is further reinforced by the retention in *Xenoturbella* of linkage groups present from sponges to vertebrates. It is interesting to note that *X. bocki* does not follow the pattern seen in other morphologically simplified animals such as nematodes and platyhelminths, which have lost and/or fused these ancestral linkage groups. We interpret this to be a signal of comparably slower genomic evolution in *Xenoturbella* in comparison to some other bilaterian lineages. The fragmented genome sequence of *Hofstenia* prevents us from asking whether the ancient linkage groups have also been preserved in the Acoelomorpha.

One of the chromosome-scale scaffolds in our assembly showed a different methylation and age signal, with both older and younger genes, and no clear relationship to metazoan linkage groups. By analyzing orthogroups of genes on this scaffold for their phylogenetic signal and finding *X. bocki* genes to cluster with those of *X. profunda* we concluded that the scaffold most likely does not represent a contamination. It remains unclear whether this scaffold is a fast-evolving chromosome, or a chromosomal fragment or arm. Very fast evolution on a chromosomal arm has for example been shown in the zebrafish^50^.

Apart from DNA from *X. bocki* we also obtained a highly contiguous genome of a species related to marine *Chlamydia* species (known from microscopy to exist in *X. bocki)*; a symbiotic relationship between *Xenoturbella* and the bacteria has been thought possible^51^. The large gene number and the completeness of genetic pathways we found in the chlamydial genome do not support an endosymbiotic relationship.

Overall, we have shown that, while *Xenoturbella* has lost some genes - in addition to the reduced number of Hox genes previously noted, we observe a reduction of some signaling pathways to the core components - in general, the *X. bocki* genome is not strikingly simpler than many other bilaterian genomes. We do not find support for a strong version of the Nephrozoa hypothesis which would predict many missing bilaterian genes. Bilaterian Hox and microRNA absent from Acoelomorpha are found in *Xenoturbella* eliminating the impact of two character types that were previously cited in support of Nephrozoa. The *Xenoturbella* genome has also largely retained the ancestral linkage groups found in other bilaterians and does not represent a structure intermediate between Eumetazoan and bilaterian ground states. Overall, while we can rule out a strong version of the Nephrozoa hypothesis with many Bilaterian characteristics missing in xenacoelomorphs, our analysis of the *Xenoturbella* genome cannot distinguish between a weak version of Nephrozoa and the Xenambulacraria topology; nevertheless, our phylogenetic analysis of gene presence and absence supports the latter.

## Methods

### Genome Sequencing, Assembly, and Scaffolding

We extracted DNA from individual *Xenoturbella* specimens with a standard and additionally worked with a Phenol-Chloroform protocol specifically developed to extract HMW DNA (dx.doi.org<u>/10.17504/protocols.io.mrxc57n</u>). The extracted DNA was quality controlled with a Nanodrop instrument in our laboratory and subsequently a TapeStation at the sequencing center. Worms were first starved and kept in repeatedly replaced salt water, reducing the likelihood of food or other contaminants in the DNA extractions. First, we sequenced Illumina short paired-end reads and mate pair libraries (see ref ^3^ for details). As the initial paired-read datasets were of low complexity and coverage, we later complemented these data with an Illumina HiSeq 2000/2500 series paired-end dataset with ∼700 bp insert size and 250bp read lengths, yielding ∼354 Million reads. Additionally, we generated ∼40 Million Illumina TruSeq Synthetic Long Reads (TSLR) for high confidence primary scaffolding.

After read cleaning with Trimmomatic v.0.38^52^ we conducted initial test assemblies using the clc assembly cell v.5 and ran the blobtools pipeline^53^ to screen for contamination (Supplementary). Not detecting any significant numbers of reads from suspicious sources in the HiSeq dataset we used SPAdes v. 3.9.0^20^ to correct and assemble a first draft genome. We also tried to use dipSPAdes but found the runtime to exceed several weeks without finishing. We submitted the SPAdes assembly to the redundans pipeline to eliminate duplicate contigs and to scaffold with all available mate pair libraries. The resulting assembly was then further scaffolded with the aid of assembled transcripts (see below) in the BADGER pipeline^54^. In this way we were able to obtain a draft genome with ∼60kb N50 that could be scaffolded to chromosome scale super-scaffolds with the use of 3C data.

We also used two remaining specimens to extract HMW DNA for Oxford Nanopore PromethION sequencing in collaboration with the Loman laboratory in Birmingham. Unfortunately, the extraction failed for one individual with the DNA appearing to be contaminated with a dark coloured residue. We were able to prepare a ligation and a PCR library for DNA from the second specimen and obtain some genomic data. However, due to pore blockage on both flow cells the combined data amounted to only about 0.5-fold coverage of the genome and was thus not useful in scaffolding. We suspect that the dark colouration of the DNA indicates a natural modification to be present in *X. bocki* DNA that inhibits sequencing with the Oxford Nanopore method.

Library preparation for genome-wide bisulfite sequencing was performed as previously described^55^. The resulting sequencing data were aligned to the *X. bocki* draft genome using Bismark in non-directional mode to identify the percentage methylation at each cytosine genome-wide. Only sites with >10 reads mapping were considered for further analysis.

### Preparation of the Hi-C libraries

The Hi-C protocol was adapted at the time from (Lieberman-Aiden et al., 2009; Sexton et al., 2012 and Marie-Nelly et al., 2014). Briefly, an animal was chemically cross-linked for one hour at room temperature in 30 mL of PBS 1X added with 3% formaldehyde (Sigma – F8775 - 4×25 mL). Formaldehyde was quenched for 20 min at RT by adding 10 ml of 2.5 M glycine. The fixed animal was recovered through centrifugation and stored at −80°C until use. To prepare the proximity ligation library, the animal was transferred to a VK05 Precellys tubes in 1X DpnII buffer (New England Biolabs; 0.5mL) and the tissues were disrupted using the Precellys Evolution homogenizer (Bertin-Instrument). SDS was added (0.3% final) to the lysate and the tubes were incubated at 65°C for 20 minutes followed by an incubation at 37°C for 30 minutes and an incubation of 30 minutes after adding 50 µL of 20% triton-X100. 150 units of the DpnII restriction enzyme were then added and the tubes were incubated overnight at 37°C. The endonuclease was inactivated 20 min at 65°C and the tubes were then centrifuged at 16,000 x g during 20 minutes, supernatant was discarded and pellets were re-suspended in 200 µl NE2 1X buffer and pooled. DNA ends were labeled using 50 µl NE2 10X buffer, 37.5 µl 0.4 mM dCTP-14-biotin, 4.5 µl 10mM dATP-dGTP-dTTP mix, 10 µl klenow 5 U/µL and incubation at 37°C for 45 minutes. The labeling mix was then transferred to ligation reaction tubes (1.6 ml ligation buffer; 160 µl ATP 100 mM; 160 µl BSA 10 mg/mL; 50 µl T4 DNA ligase (New England Biolabs, 5U/µl); 13.8 ml H2O) and incubated at 16°C for 4 hours. A proteinase K mix was added to each tube and incubated overnight at 65°C. DNA was then extracted, purified and processed for sequencing as previously described^22^. Hi-C libraries were sequenced on a NextSeq 500 (2 × 75 bp, paired-end using custom made oligonucleotides as in Marie-Nelly et al., 2014). Libraries were prepared separately on two individuals in this way but eventually merged. Note that more recent version of the HI-C protocol than the one used here have been described elsewhere^56^.

### InstaGRAAL assembly pre-processing

The primary Illumina assembly contains a number of very short contigs, which are disruptive when computing the contact distribution needed for the instaGRAAL proximity ligation scaffolding (pre-release version, see^57^ and^22^ for details). Testing several Nx metrics we found a relative length threshold, that depends on the scaffolds’ length distribution, to be a good compromise between the need for a low-noise contact distribution and the aim of connecting most of the genome. We found N90 a suitable threshold and excluded contigs below 1,308 bp. This also ensured no scaffolds shorter than three times the average length of a DpnII restriction fragment (RF) were in the assembly. In this way every contig contained enough RFs for binning and were included in the scaffolding step.

Reads from both libraries were aligned with bowtie2 (v. 2.2.5)^58^ against the DpnII RFs of the reference assembly using the hicstuff pipeline (https://github.com/koszullab/hicstuff) and in paired-end mode (with the options:-fg-maxins 5-fg-very-sensitive-local), with a mapping quality >30. The pre-processed genome was reassembled using instaGRAAL. Briefly, the program uses a Markov Chain Monte Carlo (MCMC) method that samples DNA segments (or bins) of the assembly for their best relative 1D positions with respect to each other. The quality of the positions is assessed by fitting the contact data first on a simple polymer model, then on the plot of contact frequency according to the genomic distance law computed from the data. The best relative position of a DNA segment with respect to one of its most likely neighbours consists in operations such as flips, swaps, merges or a split of contigs. Each operation is either accepted or rejected based on the computed likelihood, resulting in an iterative progression toward the 1D structure that best fits the contact data. Once the entire set of DNA segments is sampled for position (i.e. a cycle), the process starts over. The scaffolder was run independently for 50 cycles, long enough for the chromosome structure to converge. The corresponding genome is then considered stable and suitable for further analyses. The scaffolded assemblies were then refined using instaGRAAL’s instaPolish module, to correct small artefactual inversions that are sometimes a byproduct of instaGRAAL’s processing.

### Genome Annotation

#### Transcriptome Sequencing

We extracted total RNA from a single *X. bocki* individual and sequenced a strand specific Illumina paired end library. Extraction of total RNA was performed using a modified Trizol & RNeasy hybrid protocol for which tissue had to be stored in RNAlater. cDNA transcription reaction/cDNA synthesis was done using the RETROscript kit (Ambion) using both Oligo(dT) and Random Decamer primers. Detailed extraction and transcription protocols are available from the corresponding authors. The resulting transcriptomic reads (deposited under <u>SRX20415651</u>) were assembled with the Trinity pipeline^59,60^ into 103,056 sequences (N50: 705; BUSCO_v5 Eukaryota scores: C:65.1%, [S:34.1%, D:31.0%], F:22.0%, M:12.9%) for initial control and then supplied to the genome annotation pipeline (below).

### Repeat annotation

In the absence of a repeat library for Xenoturbellida we first used RepeatModeller v. 1.73 to establish a library *de novo*. We then used RepeatMasker v. 4.1.0 (https://www.repeatmasker.org) and the Dfam library^61,62^ to soft-mask the genome. We mapped the repeats to the instaGRAAL scaffolded genome with RepeatMasker.

### Gene prediction and annotation

We predicted genes using Augustus^63^ implemented into the BRAKER (v.2.1.0) pipeline^23,24^ to incorporate the RNA-Seq data. BRAKER uses spliced aligned RNA-Seq reads to improve training accuracy of the gene finder GeneMark-ET^64^. Subsequently, a highly reliable gene set predicted by GeneMark-ET in *ab initio* mode was selected to train the gene finder AUGUSTUS, which in a final step predicted genes with evidence from spliced aligned RNA-Seq reads. To make use of additional single cell transcriptome data allowing for a more precise prediction of 3’-UTRs we employed a production version of BRAKER (August 2018 snapshot). We had previously mapped the RNA-Seq data to the genome with gmap-gsnap v. 2018-07-04^65^ and used samtools^66^ and bamtools^67^ to create the necessary input files. This process was repeated in an iterative way, visually validating gene structures and comparing with mappings loci inferred from a set of single-cell RNA-Seq data (published elsewhere, see: ^68^) in particular regarding fused genes. Completeness of the gene predictions was independently assessed with BUSCO_v5^27^ setting the metazoan and the eukaryote datasets as reference respectively on gVolante^69^. We used InterProScan v. 5.27-66.0 standalone^70,71^ on the UCL cluster to annotate the predicted *X. bocki* proteins with Pfam, SUPERFAM, PANTHER, and Gene3D information.

### Horizontal Gene Transfer

To detect horizontally acquired genes in the *X. bocki* genome we used a pipeline available from (https://github.com/reubwn/hgt). Briefly, this uses blasts against the NCBI database, alignments with MAFFT^72^, and phylogenetic inferences with IQTree^73,74^ to infer most likely horizontally acquired genes, while trying to discard contamination (e.g. from co-sequenced gut microbiota).

### Orthology inference

We included 155 metazoan species and outgroups into our orthology analysis. We either downloaded available proteomes or sourced RNA-Seq reads from online repositories to then use Trinity v 2.8.5 and Trinnotate v. 3.2.0 to predict protein sets. In the latter case we implemented diamond v. 2.0.0 blast^75,76^ searches against UniProt and Pfam^77^ hmm screens against the Pfam-A dataset into the prediction process. We had initially acquired 185 datasets, but excluded some based on inferior BUSCO completeness, while at the same time aimed to span as many phyla as possible. Orthology was then inferred using Orthofinder v. 2.2.7^78,79^, again with diamond as the blast engine.

Using InterProScan v. 5.27-66.0 standalone on all proteomes we added functional annotation and then employed kinfin^33^ to summarise and analyse the orthology tables. For the kinfin analysis, we tested different query systems in regard to phylogenetic groupings (Supplementary).

To screen for inflation and contraction of gene families we first employed CAFE5^80^, but found the analysis to suffer from long branches and sparse taxon sampling in Xenambulacraria. We thus chose to query individual gene families (e.g. transcription factors) by looking up Pfam annotations in the InterProScan tables of high-quality genomes in our analysis.

Through the GenomeMaple online platform we calculated completeness of signaling pathways within the KEGG database using GhostX as the search engine.

### Presence/absence phylogenetics

We used a database of metazoan proteins, updated from ref ^81^, as the basis for an OMA analysis to calculate orthologous groups, performing two separate runs, one including *Xenoturbella* and acoels, and one with only *Xenoturbella*. We converted OMA gene OrthologousMatrix.txt files into binary gene presence absence matrices in Nexus format with datatype = restriction. We calculated phylogenetic trees on these matrices using RevBayes (see https://github.com/willpett/metazoa-gene-content for RevBayes script), as described in ref 74 with corrections for no absent sites and no singleton presence, using the reversible, not the Dollo model, as it is more likely to be able to correct for noise related to prediction errors ^82,78^. For each matrix, two runs were performed and compared and consensus trees generated with bpcomp from Phylobayes^83^.

#### Hox and ParaHox gene cluster identification and characterisation

Previous work has already used transcriptomic data and phylogenetic inference to identify the homeobox repertoire in *Xenoturbella bocki*. These annotations were used to identify genomic positions and gene annotations that correspond to Hox and ParaHox clusters in *X. bocki*. Protein sequences of homeodomains (Evx, Cdx, Gsx, antHox1, centHox1, centHox2, cent3 and postHoxP) were used as TBLASTN queries to identify putative scaffolds associated with Hox and ParaHox clusters. Gene models from these scaffolds were compared to the full length annotated homeobox transcripts from^84^ using BLASTP, using hits over 95% identity for homeobox classification. There were some possible homeodomain containing genes on the scaffolds that were not previously characterised and were therefore not given an annotation.

There were issues concerning the assignment of postHoxP and Evx to gene models. To ascertain possible CDS regions for these genes, RNA-Seq reads were mapped with HISAT2 to the scaffold and to previous annotation^84^,were assembled with Trinity and these were combined with BRAKER annotations.

Some issues were also observed with homeodomain queries matching genomic sequences that were identical, suggesting artefactual duplications. To investigate contiguity around genes the ONT reads were aligned with Minimap2 to capture long reads over regions and coverage.

### Small RNA Sequencing and Analysis

Two samples of starved worms were subjected to 5’ monophosphate dependent sequencing of RNAs between 15 and 36 nucleotides in length, according to previously described methods^85^. Using miRTrace^86^ 3.3, 18.6 million high-quality reads were extracted and merged with the 27 635 high quality 454 sequencing reads from Philippe et al. The genome sequence was screened for conserved miRNA precursors using MirMachine^87^ followed by a MirMiner run that used predicted precursors and processed and merged reads on the genome^88^. Outputs of MirMachine and MirMiner were manually curated using a uniform system for the annotation of miRNA genes^89^ and by comparing to MirGeneDB^90^.

### Neuropeptide prediction and screen

Neuropeptide prediction was conducted on the full set of *X.bocki* predicted proteins using two strategies to detect neuropeptide sequence signatures. First, using a custom script detecting the occurrence of repeated sequence patterns: RRx(3,36)RRx(3,36)RRx(3,36)RR,RRx(2,35)ZRRx(2,35)ZRR, RRx(2,35)GRRx(2,35)GRR, RRx(1,34)ZGRRx(1,34)ZGRR where R=K or R; x=any amino acid; Z=any amino acid but repeated within the pattern. Second, using HMMER3.1^91^ (hmmer.org), and a combination of neuropeptide HMM models obtained from the PFAM database (pfam.xfam.org) as well as a set of custom HMM models derived from alignment of curated sets of neuropeptide sequences^46,47,92^. Sequences retrieved using both methods and comprising fewer than 600 amino acids were further validated. First, by blast analysis: sequences with E-Value ratio “best blast hit versus ncbi nr database/best blast hit versus curated neuropeptide dataset” < 1e-40 were discarded. Second by reciprocal best blast hit clustering using Clans^93^ (eb.tuebingen.mpg.de/protein-evolution/software/clans/) with a set of curated neuropeptide sequences^46^. SignalP-5.0^94^ (cbs.dtu.dk/services/SignalP/) was used to detect the presence of a signal peptide in the curated list of predicted neuropeptide sequences while Neuropred^95^ (stagbeetle.animal.uiuc.edu/cgi-bin/neuropred.py) was used to detect cleavage sites and post-translational modifications. Sequence homology of the predicted sequence with known groups was analysed using a combination of (i) blast sequence similarity with known bilaterian neuropeptide sequences, (ii) reciprocal best blast hit clustering using Clans and sets of curated neuropeptide sequences, (iii) phylogeny using MAFFT (mafft.cbrc.jp/alignment/server/), TrimAl^96^ (trimal.cgenomics.org/) and IQ-TREE^97^ webserver for alignment, trimming and phylogeny inference respectively. Bilaterian prokineticin-like sequences were searched in ncbi nucleotide, EST and SRA databases as well as in the *Saccoglossus kowalevskii* genome assembly^74,98^ (<u>groups.oist.jp/molgenu</u>) using various bilaterian prokineticin-related protein sequences as query. Sequences used for alignments shown in figures were collected from ncbi nucleotide and protein databases as well as from the following publications: 7B2^46^; NucB2^92^; Insulin^99^; Prokineticin^37,38,100^. Alignments for figures were created with Jalview (jalview.org).

### Neuropeptide receptor search

§Neuropeptide Receptor sequences for Rhodopsin type GPCR, Secretin type GPCR and tyrosine and serine/threonine kinase receptors were searched by running HMMER3.1 on the full set of *X.bocki* predicted proteins using the 7tm_1 (PF00001), 7tm_2 (PF00002) and PK_Tyr_Ser-Thr (PF07714) HMM models respectively which were obtained from the PFAM database (pfam.xfam.org). Sequences above the significance threshold were then aligned with sequences from the curated dataset, trimmed and phylogeny inference was conducted using same method as for the neuropeptide. A second alignment and phylogeny inference was conducted after removal of all *X.bocki* sequences having no statistical support for grouping with any of the known neuropeptide receptors from the curated dataset. Curated datasets were collected from the following publications: Rhodopsin type GPCR beta and gamma and Secretin type GPCR^100^; Rhodopsin type GPCR delta (Leucine-rich repeat-containing G-protein coupled Receptors)^101^; Tyrosine kinase receptors^102,103^; and were complemented with sequences from NCBI protein database.

### Synteny

Ancestral linkage analyses rely on mutual-best-hits computed using Mmseqs2^104^ between pairs of species in which chromosomal assignments to ancestral linkage groups (ALG) was previously performed, such as *Branchiostoma floridae* or *Pecten maximus*^39^. Oxford dotplots were computed by plotting reciprocal positions of indexed pairwise orthologs between two species as performed previously^39,40^. The significance of ortholog enrichment in pairs of chromosomes was assessed using a fisher test. We also used a Python implementation of MCscanX^105^ (Haibao Tang and available on https://github.com/tanghaibao/jcvi/wiki/MCscan-(Python-version)) to compare *X. bocki* to *Euphydtia muelleri*, *Trichoplax adhearens*, *Branchiostoma floridae*, *Saccoglossus kowalevskii*, *Ciona intestinalis*, *Nematostella vectensis*, *Asteria rubens, Pecten maximus*, *Nemopilema nomurai, Carcinoscorpius rotundicauda* (see Supplementary). Briefly, the pipeline uses high quality genomes and their annotations to infer syntenic blocks based on proximity. For this an all vs. all blastp is performed and synteny extended from anchors identified in this way. Corresponding heatmaps (see Supplementary) were plotted with Python in a Jupyter notebooks instance.

### *Chlamydia* assembly and annotation

We identified a highly contiguous *Chlamydia* genome in the *X. bocki* genome assembly using blast. We then used our Oxford Nanopore derived long-reads to scaffold the *Chlamyida* genome with LINKS^106^ and annotated it with the automated PROKKA pipeline. To place the genome on the *Chlamydia* tree we extracted the 16S ribosomal RNA gene sequence, aligned it with set of *Chlamydia* 16S rRNA sequences from^28^ using MAFFT, and reconstructed the phylogeny using IQ-TREE 2^73^ We visualized the resulting tree with Figtree (http://tree.bio.ed.ac.uk/).

## Supporting information

Data Sources for Orthology

## Acknowledgements

We thank Josh Quick and Nick Loman for help with the generation of ONT long-read data. Analyses were conducted mainly on the UCL Cluster, with some computations also run on the CHEOPS cluster at the University of Cologne. We are grateful to Kevin J. Peterson for his comments on the manuscript, the miRNA section in particular. We thank the Kristineberg Center for Marine Research and Innovation for their essential support in sampling *Xenoturbella*.

## Data availability

All read sets (RNA and DNA derived) used in this study will be made available with the publication of this manuscript on the SRA database under the BioProject ID PRJNA864813. Hi-C reads are deposited under SAMN30224387, RNA-Seq under SAMN35083895. The genome assemblies of *X. bocki* (ERS12565994, ERA16814408) and the *Chlamydia* sp. (ERS12566084, ERA16814775) are deposited under PRJEB55230 at ENA.

