## Supplementary for "The slowly evolving genome of the xenacoelomorph worm *Xenoturbella bocki*"

### Supplementary information

#### HGT into the *X. bocki* genome is low

Given the close association with bacteria we were curious to see whether the *X. bocki* genome contains an elevated number of horizontally acquired genes. We did not find this to be the case. We were able to detect 56 potential horizontal gene transfer (HGT) events. Phylogenies generated using closest blast hits for each HGT candidate unveiled one of the 56 genes to be of chlamydial origin and thus likely originating from a bacterial contig. A number of the HGT candidates appear to be of Proteobacteria origin, coding for a functionally diverse set of proteins. In summary, 0.35% of the *X. bocki* genes we have identified might be horizontally acquired. See supplementary online material (at the end of this document) for alignments and gene trees.

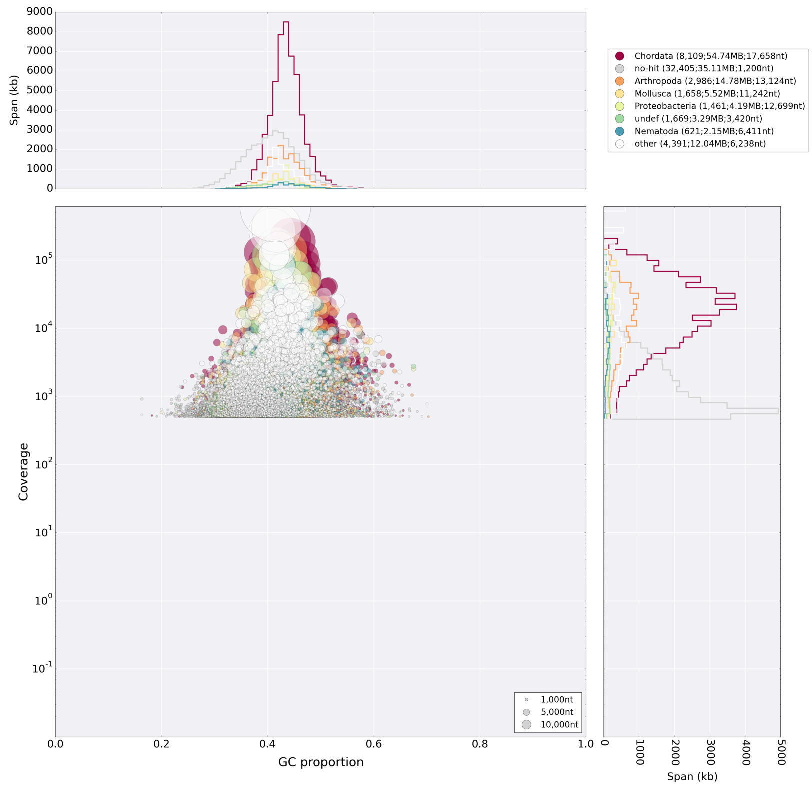


**SFigure 1.** *Blobplot analysis of the primary Illumina genome assembly.*
Performed with the SPAdes software (see Methods) the assembly shows no major microorganismal contamination, apart from the *Chlamydia* and Gammaproteobacteria described in the main text. The diamond tool was used to blast against the UniProt database for this analysis.

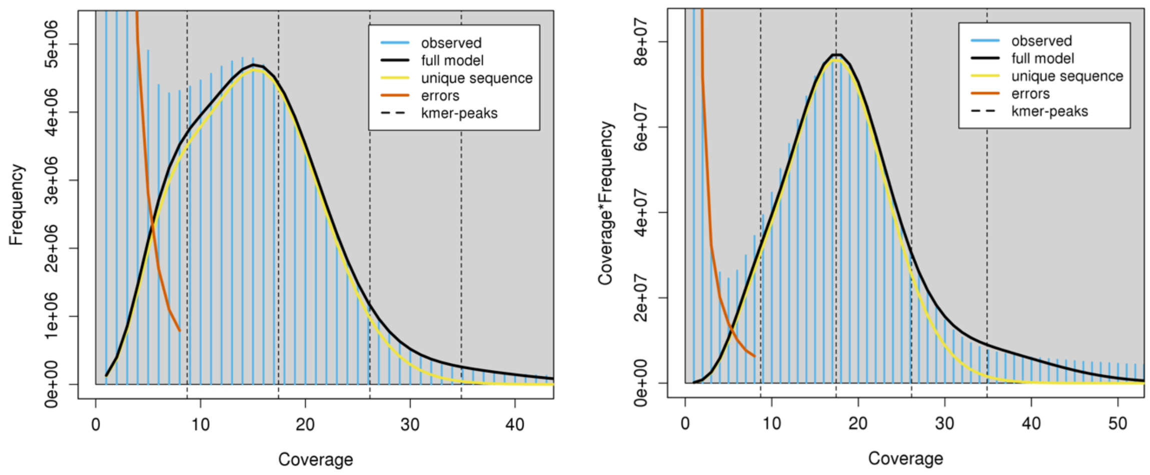


**SFigure 2.** *Kmer profile of the X. bocki Illumina WGS reads obtained with GenomeScope2 (Ranallo-Benavidez et al. 2020).* Linear plot and transformed linear plots are shown. Per description on (http://qb.cshl.edu/genomescope/) we used 21mers counted with jellyfish (Marçais and Kingsford 2011). GenomeScope genome property estimates and measures were len: 222,242,800bp, uniq: 31.7%, aa: 99.1%, ab: 0.929%, kcov: 8.72, err: 0.665%, dup: 0.527, k: 21, p:2, model fit min: 34.6%, model fit max: 96.3.

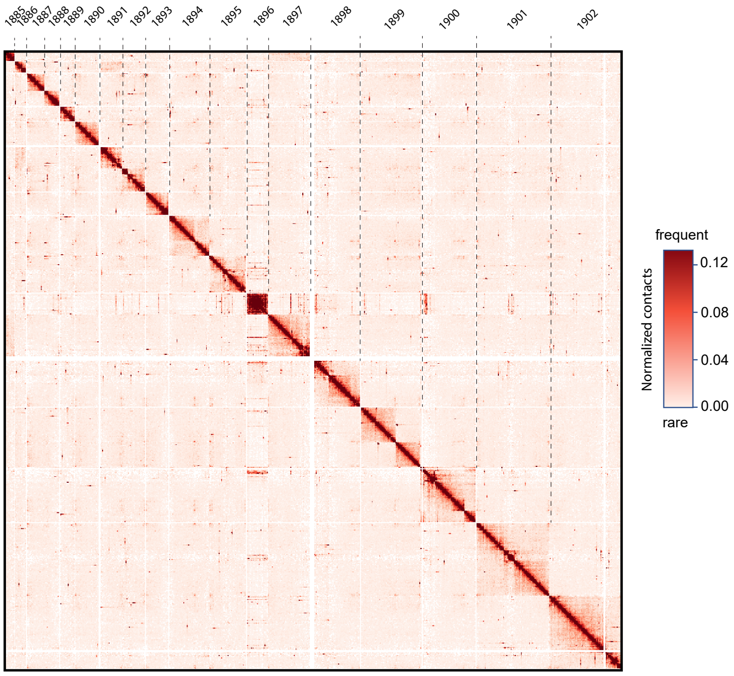


**SFigure 3.** *Contact frequency map of the largest 18 scaffolds.*

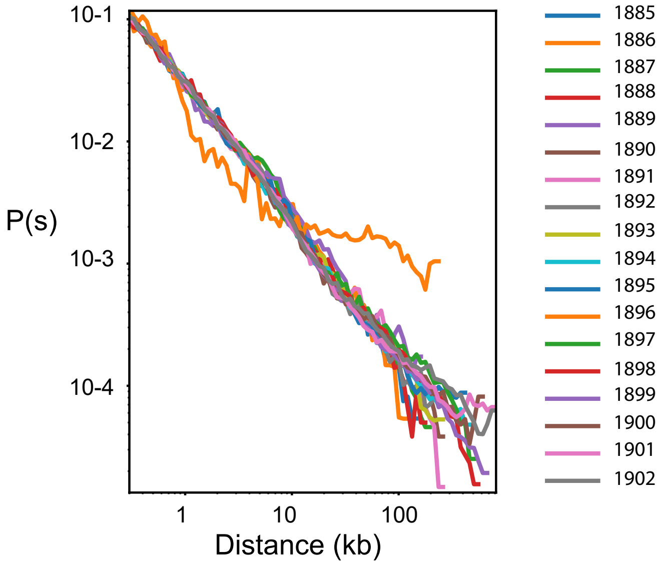


**SFigure 4.** *Distribution of contact frequency for the Hi-C data as function of distance (distance law).*

(a)

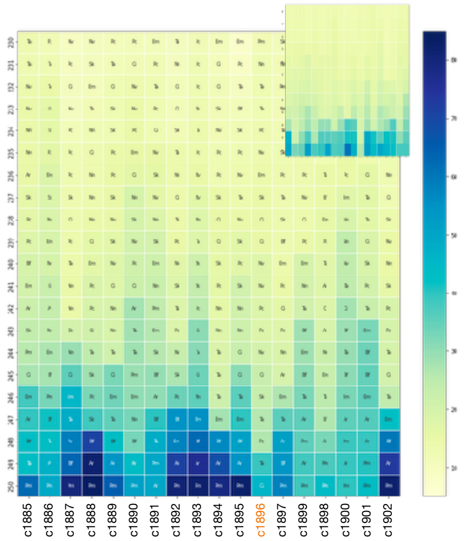


(b)

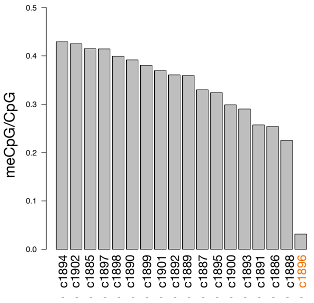

**SFigure 5.** *Conservation of metazoan synteny and Methylation in X. bocki*

(a) A summary plot of synteny between major scaffolds in the *X. bocki* genome assembly and early branching highly contiguous metazoan genome assemblies: *Euphydtia muelleri, Trichoplax adhearens, Branchiostoma floridae, Saccoglossus kowalevskii, Ciona intestinalis, Nematostella vectensis, Asteria rubens, Pecten maximus, Nemopilema nomurai, Carcinoscorpius rotundicauda*. All but one of the chromosome sized scaffolds in our assembly have at least one syntenic match in the each of the other species (see main text for one-to-one plots with key species and a description of the aberrant scaffold). We performed the same analysis with Amphioxus as the focal species as a proof of principle (inset).

(b) Analysis of methylation on the largest scaffold in the *X. bocki* genome assembly. One scaffold with a deviant gene age and synteny structure (see main text) also stands out in terms of methylation. A detailed analysis of methylation patterns across the genome and classes of genes will be published separately.

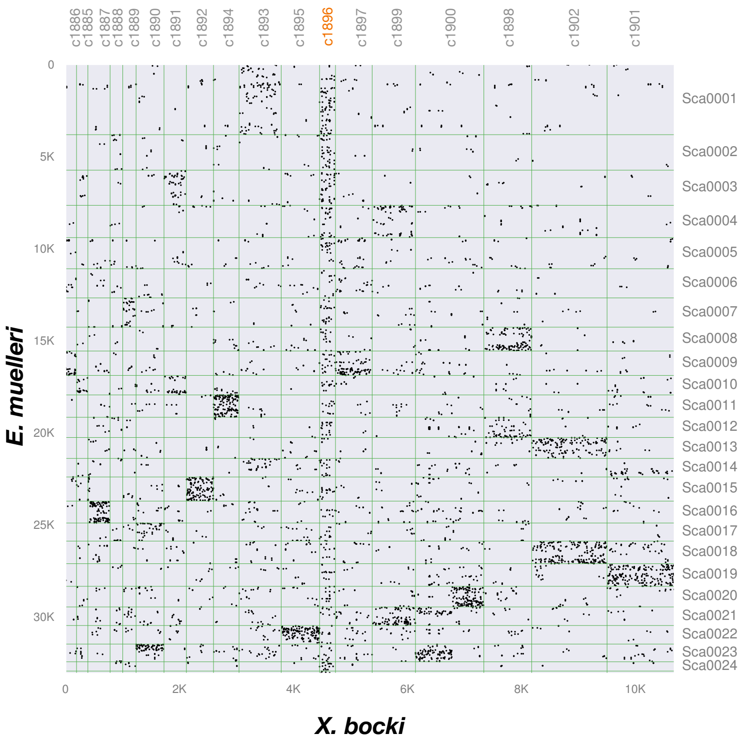


**SFigure 6.** *Intergenomic comparison of X. bocki and E. muelleri highlighting synteny connections between the aberrant scaffold c1896 and scaffolds across the sponge genome.*

#### SFigures 7 - 14: Additional information on neuropeptide signalling

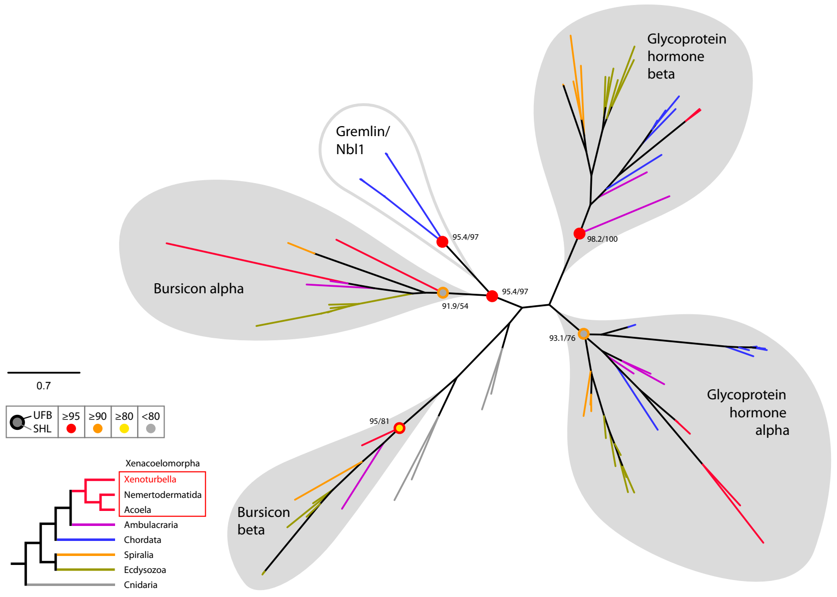


**SFigure 6**. *Radial tree representation of the phylogenetic analysis of bilaterian Glycoprotein hormone and Bursicon*. Colored dots indicate support (UFB, 1000 ultrafast bootstrap replicates; SHL, 1000 SH-aLRT replicates) and follow the color code in the left inset. Scale bar unit for branch length is the number of substitutions per site. Branches are colored according to the phylogenetic position of the organism from which the sequence originates and follow the color code in the left inset. Abbreviation: Nbl1, neuroblastoma suppressor of tumorigenicity 1 — Sequences, alignment and IQTREE tree files are available as supplementary online material.

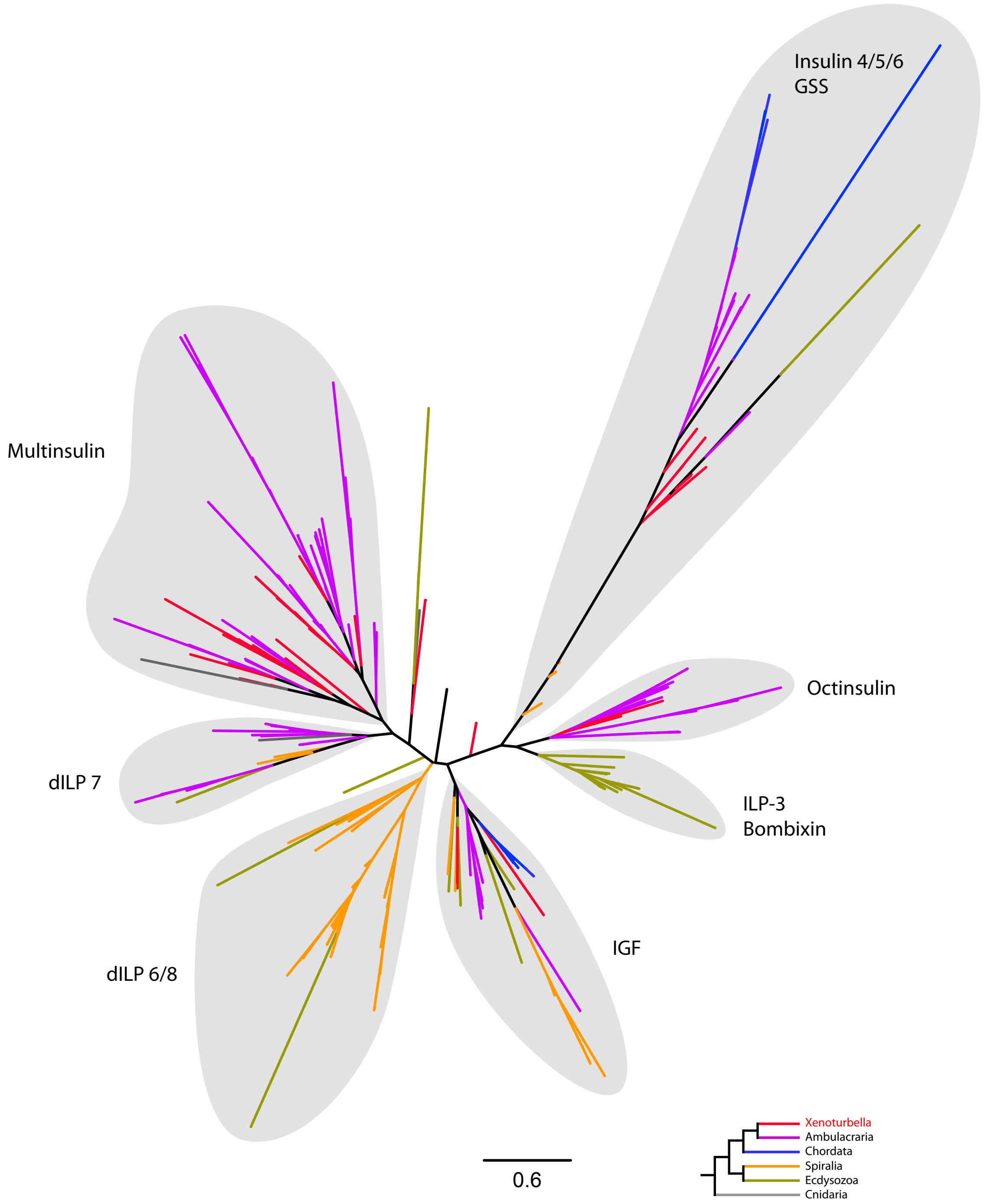
Insulin 4/5/6
GSS
Multinsulin
Octinsulin
dILP 7
ILP-3
Bombixin
IGF
dILP 6/8
0.6
Xenoturbella
Ambulacraria
Chordata
Spiralia
Ecdysozoa
Cnidaria

**SFigure 7.** *Radial tree representation of the sequence similarities analysis of bilaterian insulin related peptides.* Tree is calculated from concatenated alignment of A and B chains. Scale bar unit for branch length is the number of substitutions per site. Branches are colored according to the phylogenetic position of the organism from which the sequence originates and follow the color code in the bottom inset. Abbreviations : dILP, drosophila Insulin-like peptide ; GSS, gonad stimulating substance ; ILP, Insulin-like peptide ; IGF, Insulin-like growth factor. Sequences, alignment and IQTREE tree files are available as supplementary online material.

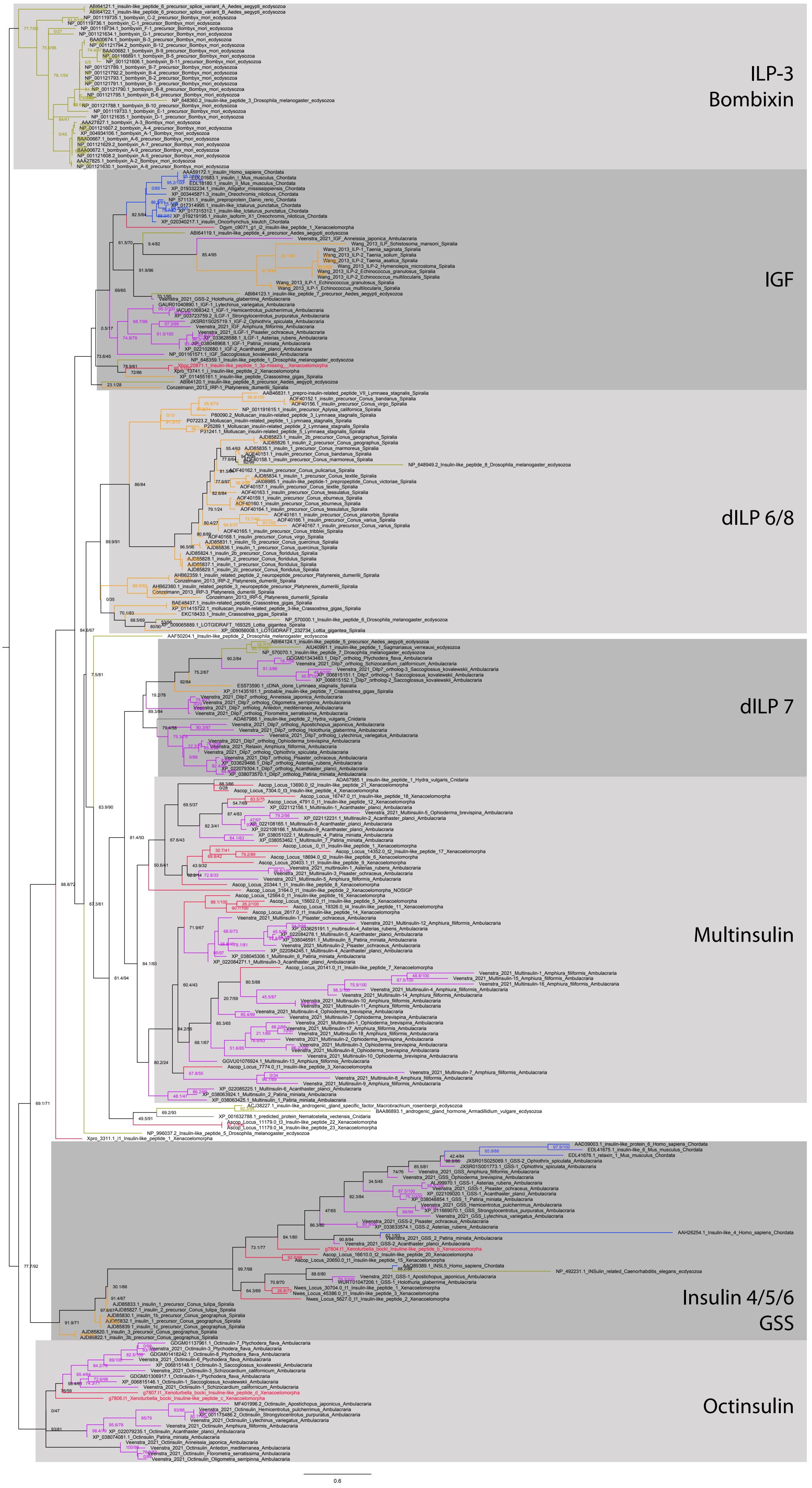
ILP-3
Bombixin
IGF
dILP 6/8
dILP 7
Multinsulin
Insulin 4/5/6
GSS
Octinsulin
0.6

**SFigure 8.** *Full tree representation of the sequence similarities analysis of bilaterian insulin related peptides.* Tree is calculated from concatenated alignment of A and B chains. Numbers represent support for nodes calculated using 1000 ultrafast bootstrap replications and 1000 SH-aLRT replicates respectively. Scale bar unit for branch length is the number of substitutions per site. Branches are colored according to the phylogenetic position of the organism from which the sequence originates : red, Xenoturbella ; pink, Ambulacraria ; blue, Chordata ; orange, Ecdysozoa ; green, Ecdysozoa ; gray, Cnidaria. Abbreviations : dILP, drosophila Insulin-like peptide ; GSS, gonad stimulating substance ; ILP, Insulin-like peptide ; IGF, Insulin-like growth factor. Sequences, alignment and IQTREE tree files are available as supplementary online material.

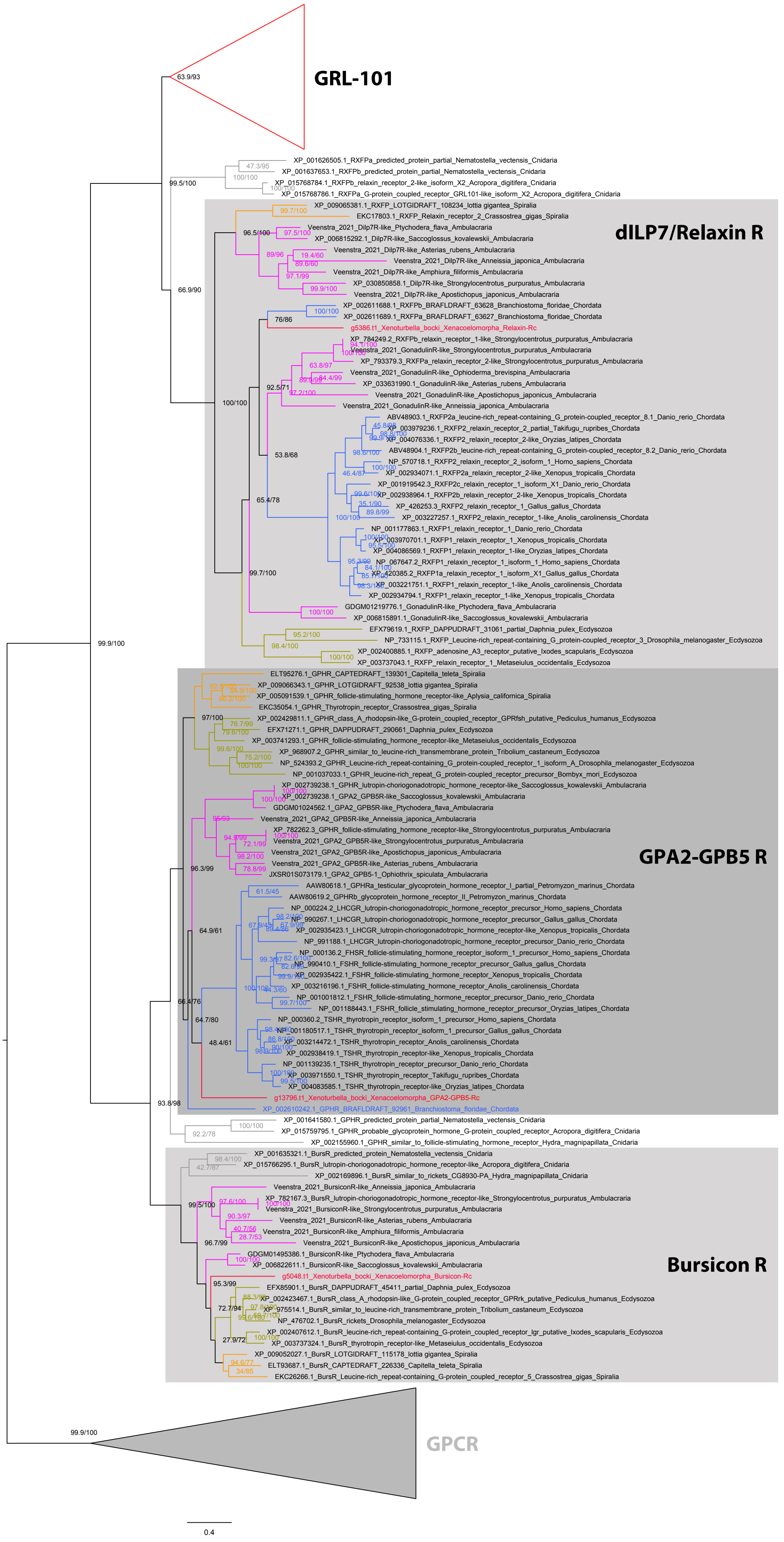

GRL-101
63.9/93
XP_001626505.1_RXFPa_predicted_protein_partial_Nematostella_vectensis_Cnidaria
XP_001637653.1_RXFPb_predicted_protein_partial_Nematostella_vectensis_Cnidaria
XP_015768784.1_RXFPb_relaxin_receptor_2-like_isoform_X2_Acropora_digitifera_Cnidaria
XP_015768786.1_RXFPa_G-protein_coupled_receptor_GRL101-like_isoform_X2_Acropora_digitifera_Cnidaria
dILP7/Relaxin R
XP_009065381.1_RXFP_LOTGIDRAFT_108234_lottia_gigantea_Spiralia
EKC17803.1_RXFP_Relaxin_receptor_2_Crassostrea_gigas_Spiralia
Veenstra_2021_Dilp7R-like_Ptychodera_flava_Ambulacraria
XP_006815292.1_Dilp7R-like_Saccoglossus_kovalewskii_Ambulacraria
Veenstra_2021_Dilp7R-like_Asterias_rubens_Ambulacraria
Veenstra_2021_Dilp7R-like_Anneissia_japonica_Ambulacraria
Veenstra_2021_Dilp7R-like_Amphiura_filiformis_Ambulacraria
XP_030850858.1_Dilp7R-like_Strongylocentrotus_purpuratus_Ambulacraria
Veenstra_2021_Dilp7R-like_Apostichopus_japonicus_Ambulacraria
XP_002611688.1_RXFPb_BRAFLDRAFT_63628_Branchiostoma_floridae_Chordata
XP_002611689.1_RXFPa_BRAFLDRAFT_63627_Branchiostoma_floridae_Chordata
g5386.t1_Xenoturbella_bocki_Xenacoelomorpha_Relaxin-Rc
XP_784249.2_RXFPb_relaxin_receptor_1-like_Strongylocentrotus_purpuratus_Ambulacraria
Veenstra_2021_GonadulinR-like_Strongylocentrotus_purpuratus_Ambulacraria
XP_793379.3_RXFPa_relaxin_receptor_2-like_Strongylocentrotus_purpuratus_Ambulacraria
Veenstra_2021_GonadulinR-like_Ophioderma_brevispina_Ambulacraria
XP_033631990.1_GonadulinR-like_Asterias_rubens_Ambulacraria
Veenstra_2021_GonadulinR-like_Apostichopus_japonicus_Ambulacraria
Veenstra_2021_GonadulinR-like_Anneissia_japonica_Ambulacraria
ABV48903.1_RXFP2a_leucine-rich_repeat-containing_G_protein-coupled_receptor_8.1_Danio_rerio_Chordata
XP_003979236.1_RXFP2_relaxin_receptor_2_partial_Takifugu_rubripes_Chordata
XP_004076336.1_RXFP2_relaxin_receptor_2-like_Oryzias_latipes_Chordata
ABV48904.1_RXFP2b_leucine-rich_repeat-containing_G_protein-coupled_receptor_8.2_Danio_rerio_Chordata
NP_570718.1_RXFP2_relaxin_receptor_2_isoform_1_Homo_sapiens_Chordata
XP_002934071.1_RXFP2a_relaxin_receptor_2-like_Xenopus_tropicalis_Chordata
XP_001919542.3_RXFP2c_relaxin_receptor_1_isoform_X1_Danio_rerio_Chordata
XP_002938964.1_RXFP2b_relaxin_receptor_2-like_Xenopus_tropicalis_Chordata
XP_426253.3_RXFP2_relaxin_receptor_2_Gallus_gallus_Chordata
XP_003227257.1_RXFP2_relaxin_receptor_1-like_Anolis_carolinensis_Chordata
NP_001177863.1_RXFP1_relaxin_receptor_1_Danio_rerio_Chordata
XP_003970701.1_RXFP1_relaxin_receptor_1_Xenopus_tropicalis_Chordata
XP_004086569.1_RXFP1_relaxin_receptor_1-like_Oryzias_latipes_Chordata
NP_067647.2_RXFP1_relaxin_receptor_1_isoform_1_Homo_sapiens_Chordata
XP_420385.2_RXFP1a_relaxin_receptor_1_isoform_X1_Gallus_gallus_Chordata
XP_003221751.1_RXFP1_relaxin_receptor_1-like_Anolis_carolinensis_Chordata
XP_002934794.1_RXFP1_relaxin_receptor_1-like_Xenopus_tropicalis_Chordata
GDGM01219776.1_GonadulinR-like_Ptychodera_flava_Ambulacraria
XP_006815891.1_GonadulinR-like_Saccoglossus_kovalewskii_Ambulacraria
EFX79619.1_RXFP_DAPPUDRAFT_31061_partial_Daphnia_pulex_Ecdysozoa
NP_733115.1_RXFP_Leucine-rich_repeat-containing_G_protein-coupled_receptor_3_Drosophila_melanogaster_Ecdysozoa
XP_002400885.1_RXFP_adenosine_A3_receptor_putative_Ixodes_scapularis_Ecdysozoa
XP_003737043.1_RXFP_relaxin_receptor_1_Metaseiulus_occidentalis_Ecdysozoa
GPA2-GPB5 R
ELT95276.1_GPHR_CAPTEDRAFT_139301_Capitella_teleta_Spiralia
XP_009066343.1_GPHR_LOTGIDRAFT_92538_lottia_gigantea_Spiralia
XP_005091539.1_GPHR_follicle-stimulating_hormone_receptor-like_Aplysia_californica_Spiralia
EKC35054.1_GPHR_Thyrotropin_receptor_Crassostrea_gigas_Spiralia
XP_002429811.1_GPHR_class_A_rhodopsin-like_G-protein_coupled_receptor_GPRfsh_putative_Pediculus_humanus_Ecdysozoa
EFX71271.1_GPHR_DAPPUDRAFT_290661_Daphnia_pulex_Ecdysozoa
XP_003741293.1_GPHR_follicle-stimulating_hormone_receptor-like_Metaseiulus_occidentalis_Ecdysozoa
XP_968907.2_GPHR_similar_to_leucine-rich_transmembrane_protein_Tribolium_castaneum_Ecdysozoa
NP_524393.2_GPHR_Leucine-rich_repeat-containing_G_protein-coupled_receptor_1_isoform_A_Drosophila_melanogaster_Ecdysozoa
NP_001037033.1_GPHR_leucine-rich_repeat_G_protein-coupled_receptor_precursor_Bombyx_mori_Ecdysozoa
XP_002739238.1_GPHR_lutropin-choriogonadotropic_hormone_receptor-like_Saccoglossus_kowalevskii_Ambulacraria
XP_002739238.1_GPA2_GPB5R-like_Saccoglossus_kowalevskii_Ambulacraria
GDGM01024562.1_GPA2_GPB5R-like_Ptychodera_flava_Ambulacraria
Veenstra_2021_GPA2_GPB5R-like_Anneissia_japonica_Ambulacraria
XP_782262.3_GPHR_follicle-stimulating_hormone-like_Strongylocentrotus_purpuratus_Ambulacraria
Veenstra_2021_GPA2_GPB5R-like_Strongylocentrotus_purpuratus_Ambulacraria
Veenstra_2021_GPA2_GPB5R-like_Apostichopus_japonicus_Ambulacraria
Veenstra_2021_GPA2_GPB5R-like_Asterias_rubens_Ambulacraria
JXSR01S073179.1_GPA2_GPB5-1_Ophiothrix_spiculata_Ambulacraria
AAW80618.1_GPHRa_testicular_glycoprotein_hormone_receptor_I_partial_Petromyzon_marinus_Chordata
AAW80619.2_GPHRb_glycoprotein_hormone_receptor_II_Petromyzon_marinus_Chordata
NP_000224.2_LHCGR_lutropin-choriogonadotropic_hormone_receptor_precursor_Homo_sapiens_Chordata
NP_990267.1_LHCGR_lutropin-choriogonadotropic_hormone_receptor_precursor_Gallus_gallus_Chordata
XP_002935423.1_LHCGR_lutropin-choriogonadotropic_hormone_receptor-like_Xenopus_tropicalis_Chordata
NP_991188.1_LHCGR_lutropin-choriogonadotropic_hormone_receptor_precursor_Danio_rerio_Chordata
NP_000136.2_FHSR_follicle-stimulating_hormone_receptor_isoform_1_precursor_Homo_sapiens_Chordata
NP_990410.1_FSHR_follicle-stimulating_hormone_receptor_precursor_Gallus_gallus_Chordata
XP_002935422.1_FSHR_follicle-stimulating_hormone_receptor_Xenopus_tropicalis_Chordata
XP_003216196.1_FSHR_follicle-stimulating_hormone_receptor_Anolis_carolinensis_Chordata
NP_001001812.1_FSHR_follicle-stimulating_hormone_receptor_precursor_Danio_rerio_Chordata
NP_001188443.1_FSHR_follicle_stimulating_hormone_receptor_precursor_Oryzias_latipes_Chordata
NP_000360.2_TSHR_thyrotropin_receptor_isoform_1_precursor_Homo_sapiens_Chordata
NP_001180517.1_TSHR_thyrotropin_receptor_isoform_1_precursor_Gallus_gallus_Chordata
XP_003214472.1_TSHR_thyrotropin_receptor_Anolis_carolinensis_Chordata
XP_002938419.1_TSHR_thyrotropin_receptor-like_Xenopus_tropicalis_Chordata
NP_001139235.1_TSHR_thyrotropin_receptor_precursor_Danio_rerio_Chordata
XP_003971550.1_TSHR_thyrotropin_receptor_Takifugu_rubripes_Chordata
XP_004083585.1_TSHR_thyrotropin_receptor-like_Oryzias_latipes_Chordata
g13796.t1_Xenoturbella_bocki_Xenacoelomorpha_GPA2-GPB5-Rc
XP_002610242.1_GPHR_BRAFLDRAFT_92961_Branchiostoma_floridae_Chordata
XP_001641580.1_GPHR_predicted_protein_partial_Nematostella_vectensis_Cnidaria
XP_015759795.1_GPHR_probable_glycoprotein_hormone_G-protein_coupled_receptor_Acropora_digitifera_Cnidaria
XP_002155960.1_GPHR_similar_to_follicle-stimulating_hormone_receptor_Hydra_magnipapillata_Cnidaria
XP_001635321.1_BursR_predicted_protein_Nematostella_vectensis_Cnidaria
XP_015766295.1_BursR_lutropin-choriogonadotropic_hormone_receptor-like_Acropora_digitifera_Cnidaria
XP_002169896.1_BursR_similar_to_rickets_CG8930-PA_Hydra_magnipapillata_Cnidaria
Veenstra_2021_BursiconR-like_Anneissia_japonica_Ambulacraria
XP_782167.3_BursR_lutropin-choriogonadotropic_hormone_receptor-like_Strongylocentrotus_purpuratus_Ambulacraria
Veenstra_2021_BursiconR-like_Strongylocentrotus_purpuratus_Ambulacraria
Veenstra_2021_BursiconR-like_Asterias_rubens_Ambulacraria
Veenstra_2021_BursiconR-like_Amphiura_filiformis_Ambulacraria
Veenstra_2021_BursiconR-like_Apostichopus_japonicus_Ambulacraria
GDGM01495386.1_BursiconR-like_Ptychodera_flava_Ambulacraria
XP_006822611.1_BursiconR-like_Saccoglossus_kovalewskii_Ambulacraria
g5048.t1_Xenoturbella_bocki_Xenacoelomorpha_Bursicon-Rc
EFX85901.1_BursR_DAPPUDRAFT_45411_partial_Daphnia_pulex_Ecdysozoa
XP_002423467.1_BursR_class_A_rhodopsin-like_G-protein_coupled_receptor_GPRrk_putative_Pediculus_humanus_Ecdysozoa
XP_975514.1_BursR_similar_to_leucine-rich_transmembrane_protein_Tribolium_castaneum_Ecdysozoa
NP_476702.1_BursR_rickets_Drosophila_melanogaster_Ecdysozoa
XP_002407612.1_BursR_leucine-rich_repeat-containing_G-protein_coupled_receptor_lgr_putative_Ixodes_scapularis_Ecdysozoa
XP_003737324.1_BursR_thyrotropin_receptor-like_Metaseiulus_occidentalis_Ecdysozoa
XP_009052027.1_BursR_LOTGIDRAFT_115178_lottia_gigantea_Spiralia
ELT93687.1_BursR_CAPTEDRAFT_226336_Capitella_teleta_Spiralia
EKC26266.1_BursR_Leucine-rich_repeat-containing_G-protein_coupled_receptor_5_Crassostrea_gigas_Spiralia
Bursicon R
99.9/100
GPCR
0.4

**SFigure 9.** *Full tree representation of the phylogenetic analysis of bilaterian Leucine-rich repeat-containing G-protein coupled Receptors (Rhodopsin type G-protein coupled Receptors delta).* Numbers represent support for nodes calculated using 1000 Ultrafast bootstrap replications and 1000 SH-aLRT replicates respectively. Scale bar unit for branch length is the number of substitutions per site. Branches are colored according to the phylogenetic position of the organism from which the sequence originates : red, Xenoturbella ; pink, Ambulacraria ; blue, Chordata ; orange, Ecdysozoa ; green, Ecdysozoa ; gray, Cnidaria. Collapsed group colored in red indicate that they contain at least one *Xenoturbella bocki* sequence. Abbreviations : GPA2, Glycoprotein Hormone alpha5 ; GPB5, Glycoprotein Hormone beta2 ; GPCR, G Protein-Coupled Receptor ; GRL-101, G-protein coupled receptor GRL101. Sequences, alignment and IQTREE tree files are available as supplementary online material.

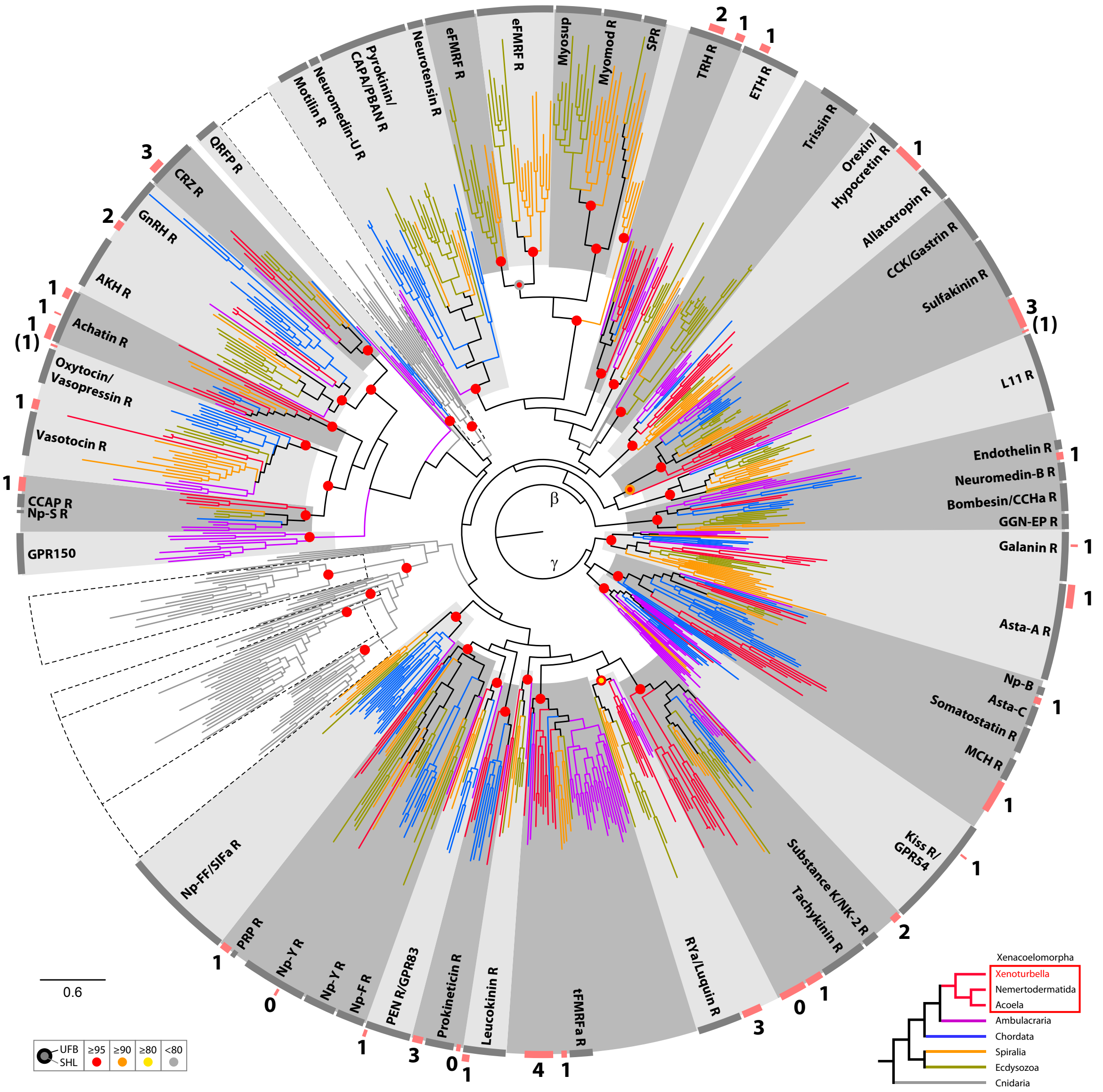

**SFigure 10.** *Circular tree representation of the phylogenetic analysis of bilaterian Rhodopsin type G-protein coupled Receptors beta and gamma.* Colored dots indicate support (UFB, 1000 ultrafast bootstrap replicates; SHL, 1000 SH-aLRT replicates) for main nodes and

follow the color code in the bottom inset. Scale bar unit for branch length is the number of substitutions per site. Branches are colored according to the phylogenetic position of the organism from which the sequence originates and follow the color code in the bottom inset. Circular gray bars highlights names of groups of annotated sequences. Circular red bars indicate position of groups of Xenacoelomorpha sequences and associated number the number of Xenoturbella bocki sequence(s) within these groups. Abbreviations : AKH, adipokinetic hormone ; Asta-A, Allatostatin-A ; Asta-C, Allatostatin-C ; CAPA, Cardio acceleratory peptide ; CCAP, crustacean cardioactive peptide ; CCHa, CCHamide peptide ; CCK, cholecystokinin ; CRZ, Corazonin ; eFMRF, ecdysozoan-FMRFamide peptide ; GGN-EP, GGN excitatory peptide ; ETH, ecdysis triggering hormone ; GnRH, Gonadotropin Releasing Hormone ; GPR150, G Protein-Coupled Receptor 150 ; GPR54, G Protein-Coupled Receptor 54 ; GPR83, G Protein-Coupled Receptor 83 ; MCH, melanin concentrating hormone ; Myomod, Myomodulin ; NK-2, Neurokinin 2 ; Np-B/W, Neuropeptide B/W ; Np-FF, Neuropeptide FF ; Np-F, Neuropeptide F ; Np-S, Neuropeptide S ; Np-Y, Neuropeptide Y ; PBAN, pheromone biosynthesis activation neuropeptide ; PEN, neuroendocrine peptide PEN ; PRP, Prolactin releasing peptide ; QRFP, Neuropeptide QRFP ; RYa, RYamide peptide ; SIFa, SIFamide peptide ; SPR, Sex peptide receptor ; tFMRFa, trochozoan-FMRFamide peptide ; TRH, thyrotrophin-releasing hormone. Sequences, alignment and IQTREE tree files are available as supplementary online material.

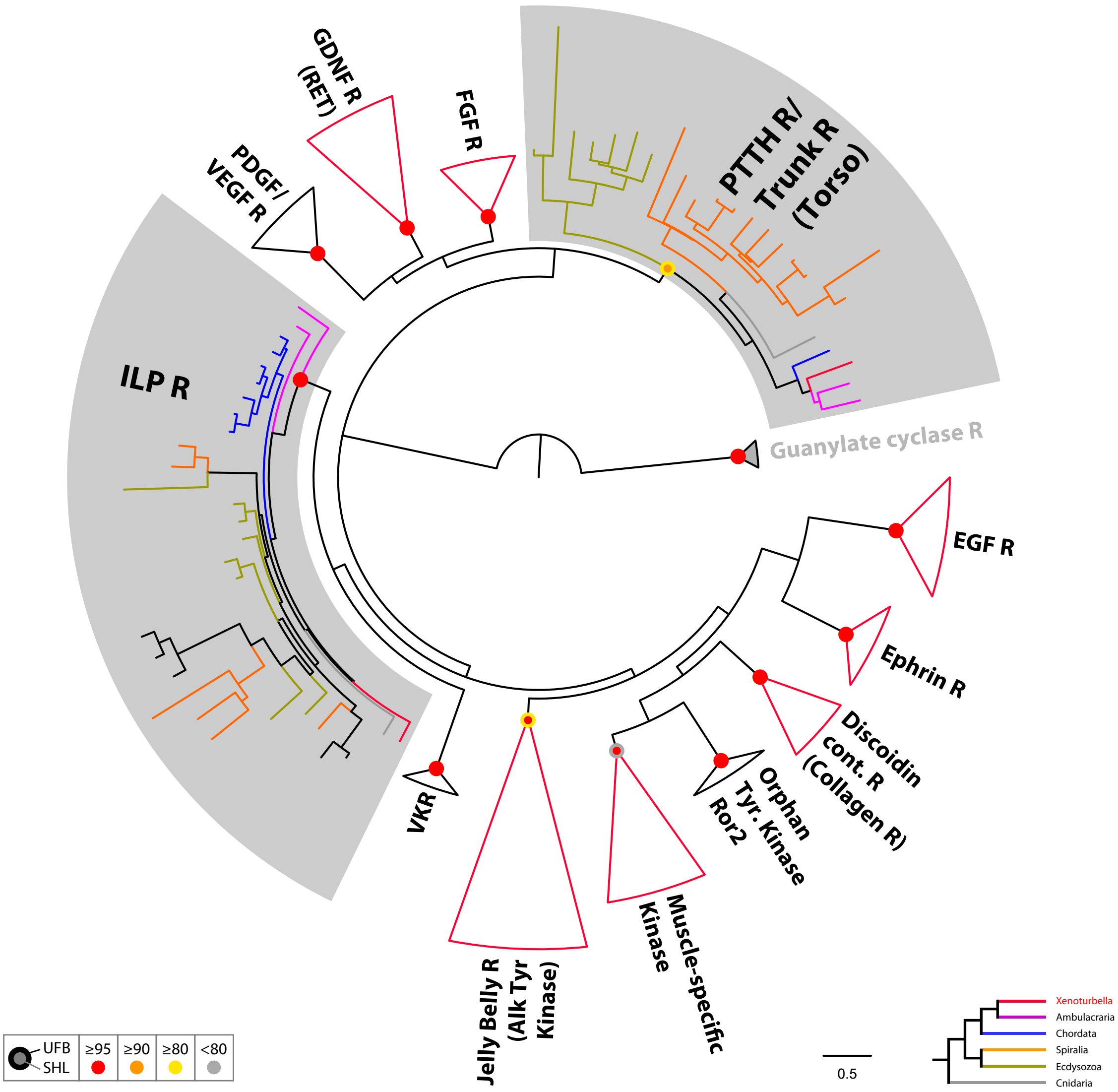


**SFigure 11.** *Circular tree representation of the phylogenetic analysis of bilaterian Tyrosine kinase Receptors.* Colored dots indicate support (UFB, 1000 ultrafast bootstrap

replicates; SHL, 1000 SH-aLRT replicates) and follow the color code in the bottom inset. Scale bar unit for branch length is the number of substitutions per site. Branches are colored according to the phylogenetic position of the organism from which the sequence originates and follow the color code in the bottom inset. Collapsed group colored in red indicate that they contain at least one *Xenoturbella bocki* sequence. Abbreviations : EGF, Epidermal Growth Factor ; Discoidin cont. R, discoidin domain-containing receptor ; Orphan Tyr. Kinase Ror2, receptor tyrosine kinase-like orphan receptor 2 ; VKR, Venus kinase Receptor ; ILP, Insulin-like peptide ; PDGF, Platelet-derived growth factor ; VEGF, Vascular endothelial growth factor ; GDNF, Glial cell line-derived neurotrophic factor ; FGF, fibroblast growth factor ; PTTH, Prothoracicotropic hormone. Sequences, alignment and IQTREE tree files are available as supplementary online material.

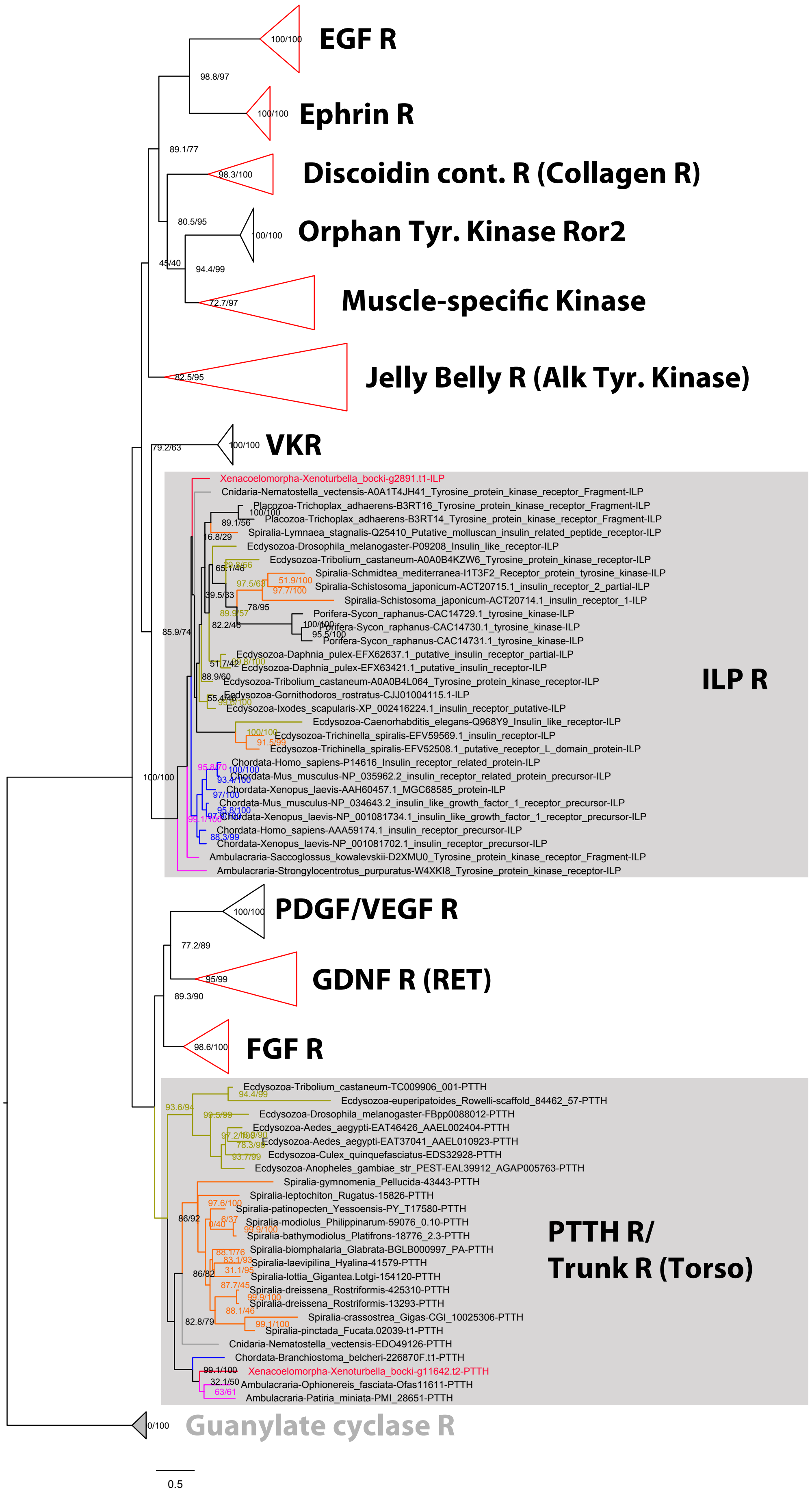
EGF R
Ephrin R
Discoidin cont. R (Collagen R)
Orphan Tyr. Kinase Ror2
Muscle-specific Kinase
Jelly Belly R (Alk Tyr. Kinase)
VKR
Xenacoelomorpha-Xenoturbella_bocki-g2891.t1-ILP
Cnidaria-Nematostella_vectensis-A0A1T4JH41_Tyrosine_protein_kinase_receptor_Fragment-ILP
Placozoa-Trichoplax_adhaerens-B3RT16_Tyrosine_protein_kinase_receptor_Fragment-ILP
Placozoa-Trichoplax_adhaerens-B3RT14_Tyrosine_protein_kinase_receptor_Fragment-ILP
Spiralia-Lymnaea_stagnalis-Q25410_Putative_molluscan_insulin_related_peptide_receptor-ILP
Ecdysozoa-Drosophila_melanogaster-P09208_Insulin_like_receptor-ILP
Ecdysozoa-Tribolium_castaneum-A0A0B4KZW6_Tyrosine_protein_kinase_receptor-ILP
Spiralia-Schmidtea_mediterranea-I1T3F2_Receptor_protein_tyrosine_kinase-ILP
Spiralia-Schistosoma_japonicum-ACT20715.1_insulin_receptor_2_partial-ILP
Spiralia-Schistosoma_japonicum-ACT20714.1_insulin_receptor_1-ILP
Porifera-Sycon_raphanus-CAC14729.1_tyrosine_kinase-ILP
Porifera-Sycon_raphanus-CAC14730.1_tyrosine_kinase-ILP
Porifera-Sycon_raphanus-CAC14731.1_tyrosine_kinase-ILP
Ecdysozoa-Daphnia_pulex-EFX62637.1_putative_insulin_receptor_partial-ILP
Ecdysozoa-Daphnia_pulex-EFX63421.1_putative_insulin_receptor-ILP
Ecdysozoa-Tribolium_castaneum-A0A0B4L064_Tyrosine_protein_kinase_receptor-ILP
Ecdysozoa-Gornithodoros_rostratus-CJJ01004115.1-ILP
Ecdysozoa-Ixodes_scapularis-XP_002416224.1_insulin_receptor_putative-ILP
Ecdysozoa-Caenorhabditis_elegans-Q968Y9_Insulin_like_receptor-ILP
Ecdysozoa-Trichinella_spiralis-EFV59569.1_insulin_receptor-ILP
Ecdysozoa-Trichinella_spiralis-EFV52508.1_putative_receptor_L_domain_protein-ILP
Chordata-Homo_sapiens-P14616_Insulin_receptor_related_protein-ILP
Chordata-Mus_musculus-NP_035962.2_insulin_receptor_related_protein_precursor-ILP
Chordata-Xenopus_laevis-AAH60457.1_MGC68585_protein-ILP
Chordata-Mus_musculus-NP_034643.2_insulin_like_growth_factor_1_receptor_precursor-ILP
Chordata-Xenopus_laevis-NP_001081734.1_insulin_like_growth_factor_1_receptor_precursor-ILP
Chordata-Homo_sapiens-AAA59174.1_insulin_receptor_precursor-ILP
Chordata-Xenopus_laevis-NP_001081702.1_insulin_receptor_precursor-ILP
Ambulacraria-Saccoglossus_kowalevskii-D2XMU0_Tyrosine_protein_kinase_receptor_Fragment-ILP
Ambulacraria-Strongylocentrotus_purpuratus-W4XKI8_Tyrosine_protein_kinase_receptor-ILP
ILP R
PDGF/VEGF R
GDNF R (RET)
FGF R
Ecdysozoa-Tribolium_castaneum-TC009906_001-PTTH
Ecdysozoa-euperipatoides_Rowelli-scaffold_84462_57-PTTH
Ecdysozoa-Drosophila_melanogaster-FBpp0088012-PTTH
Ecdysozoa-Aedes_aegypti-EAT46426_AAEL002404-PTTH
Ecdysozoa-Aedes_aegypti-EAT37041_AAEL010923-PTTH
Ecdysozoa-Culex_quinquefasciatus-EDS32928-PTTH
Ecdysozoa-Anopheles_gambiae_str_PEST-EAL39912_AGAP005763-PTTH
Spiralia-gymnomenia_Pellucida-43443-PTTH
Spiralia-leptochiton_Rugatus-15826-PTTH
Spiralia-patinopecten_Yessoensis-PY_T17580-PTTH
Spiralia-modiolus_Philippinarum-59076_0.10-PTTH
Spiralia-bathymodiolus_Platifrons-18776_2.3-PTTH
Spiralia-biomphalaria_Glabrata-BGLB000997_PA-PTTH
Spiralia-laevipilina_Hyalina-41579-PTTH
Spiralia-lottia_Gigantea.Lotgi-154120-PTTH
Spiralia-dreissena_Rostriformis-425310-PTTH
Spiralia-dreissena_Rostriformis-13293-PTTH
Spiralia-crassostrea_Gigas-CGI_10025306-PTTH
Spiralia-pinctada_Fucata.02039-t1-PTTH
Cnidaria-Nematostella_vectensis-EDO49126-PTTH
Chordata-Branchiostoma_belcheri-226870F.t1-PTTH
Xenacoelomorpha-Xenoturbella_bocki-g11642.t2-PTTH
Ambulacraria-Ophionereis_fasciata-Ofas11611-PTTH
Ambulacraria-Patiria_miniata-PMI_28651-PTTH
PTTH R/
Trunk R (Torso)
Guanylate cyclase R
0.5

**SFigure 12.** *Full tree representation of the phylogenetic analysis of bilaterian Tyrosine kinase Receptors.* Numbers represent support for nodes calculated using 1000 ultrafast bootstrap replications and 1000 SH-aLRT replicates respectively. Scale bar unit for branch length is the number of substitutions per site. Branches are colored according to the phylogenetic position of the organism from which the sequence originates : red, Xenoturbella ; pink, Ambulacraria ; blue, Chordata ; orange, Ecdysozoa ; green, Ecdysozoa ; gray, Cnidaria. Collapsed group colored in red indicate that they contain at least one *Xenoturbella bocki* sequence. Abbreviations : EGF, Epidermal Growth Factor ; Discoidin cont. R, discoidin domain-containing receptor ; Orphan Tyr. Kinase Ror2, receptor tyrosine kinase-like orphan receptor 2 ; VKR, Venus kinase Receptor ; ILP, Insulin-like peptide ; PDGF, Platelet-derived growth factor ; VEGF, Vascular endothelial growth factor ; GDNF, Glial cell line-derived neurotrophic factor ; FGF, fibroblast growth factor ; PTTH, Prothoracicotropic. Sequences, alignment and IQTREE tree files are available as supplementary online material.

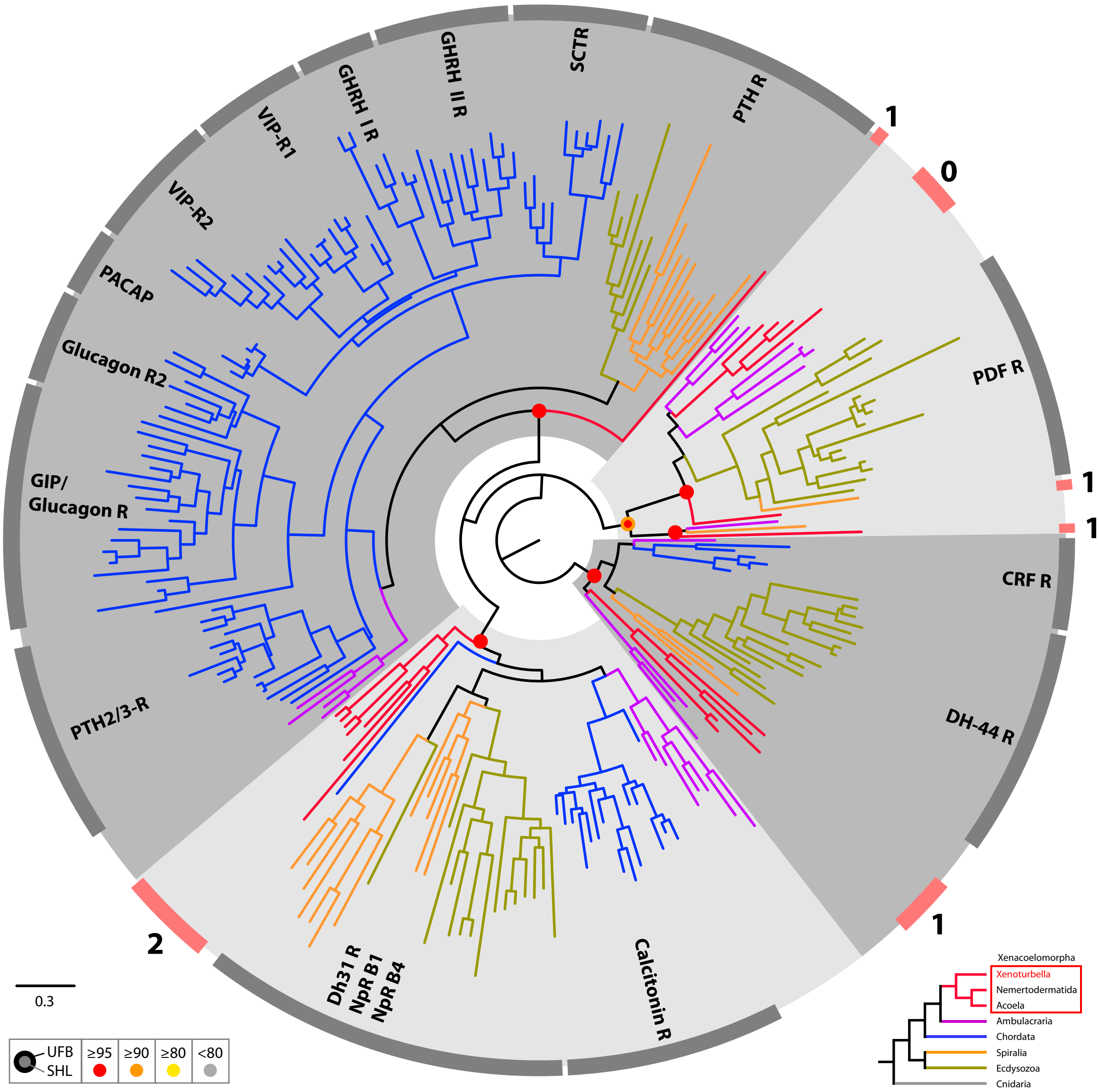
PTH R
SCTR
GHRH II R
GHRH I R
VIP-R1
VIP-R2
PACAP
Glucagon R2
GIP/
Glucagon R
PTH2/3-R
Dh31 R
NpR B1
NpR B4
Calcitonin R
DH-44 R
CRF R
PDF R
0
1
1
1
1
2
0.3
UFB
SHL
≥95
≥90
≥80
<80
Xenacoelomorpha
Xenoturbella
Nemertodermatida
Acoela
Ambulacraria
Chordata
Spiralia
Ecdysozoa
Cnidaria

**SFigure 13.** *Circular tree representation of the phylogenetic analysis of bilaterian Secretin type G-protein coupled Receptors.* Colored dots indicate support (UFB, 1000 ultrafast bootstrap replicates; SHL, 1000 SH-aLRT replicates) for main nodes and follow the color code in the bottom inset. Scale bar unit for branch length is the number of substitutions per site. Branches are colored according to the phylogenetic position of the organism from which the sequence originates and follow the color code in the bottom inset. Circular gray bars highlights names of groups of annotated sequences. Circular red bars indicate position of groups of Xenacoelomorpha sequences and associated number the number of Xenoturbella bocki sequence(s) within these groups. Abbreviations : DH31, diuretic hormone 31 ; Np-R B1, Neuropeptide receptor B3 ; Np-R B4, Neuropeptide receptor B1 ; PDF, Pigment-dispersing factor ; CRF, Corticotropin-releasing factor ; DH-44, diuretic hormone 44 ; PTH2/3-R, Parathyroid hormone receptor2/3 ; GIP, Gastric inhibitory polypeptide ; PACAP, Pituitary adenylate cyclase-activating polypeptide ; VIP-R, Vasoactive intestinal polypeptide receptor ; GHRH, Growth hormone-releasing hormone ; PTH, Parathyroid hormone receptor ; SCTR, Secretin Receptor. Sequences, alignment and IQTREE tree files are available as supplementary online material.

**Supplementary Table 1.** *Improvement of assembly and scaffolding metrics.*

| Assembly step | # seqs | # reals | # Ns | Max length | N50 |
|---|---|---|---|---|---|
| redundans contigs | 37,880 | 113,212,556 | 38,3327 | 206,709 | 8,544 |
| redundans scaffolds | 24,538 | 117,405,089 | 3,021,351 | 952,321 | 52,073 |
| pre instaGRAAL | 23,094 | 117,396,873 | 3,534,582 | 960,978 | 61,989 |
| final scaffolds | 27,939 | 107,712,917 | 3,328,069 | 8,757,424 | 2,730,651 |

Assessed with the jvci toolbox: https://github.com/tanghaibao/jcvi.git.

**Supplementary online material on Zenodo (doi:10.5281/zenodo.6962271):**

S. File 1: Orthofinder Orthology, Xbocki_genome_Orthogroups.csv.gz;
S. File 2: Comparative annotation of KEGG pathway completeness, Xbocki_genome.KEGG_module_completeness.xls;
S. File 3: Neuropetide screen, alignments and treefiles;
S. File 4: 16S rRNA tree for Chlamydia species, Chlamydia_16S_rRNA.tree;
S. File 5: Alignments and trees from HGT screen, HGT_screen_aln-tree.tgz.
S. File 6: Excel table with data sources for OrthoFinder analysis.
